# Miniature inverted-repeat transposable elements drive rapid microRNA diversification in angiosperms

**DOI:** 10.1101/2021.07.25.453727

**Authors:** Zhonglong Guo, Zheng Kuang, Yihan Tao, Haotian Wang, Miaomiao Wan, Chen Hao, Fei Shen, Xiaozeng Yang, Lei Li

## Abstract

MicroRNAs (miRNAs) are rapidly evolving endogenous small RNAs programing organism function and behavior. Although models for miRNA origination have been proposed based on sporadic cases, the genomic mechanisms driving swift diversification of the miRNA repertoires in plants remain elusive. Here, by comprehensively analyzing 20 phylogenetically representative plant species, we identified miniature inverted-repeat transposable elements (MITEs) as the predominant genomic sources for *de novo* miRNAs in angiosperms. Our data illustrated a transposition-transcription process whereby properly sized MITEs transposed into active genic regions could be converted into new miRNAs, termed MITE-miRNAs, in as few as 20 generations. We showed that this molecular domestication mechanism leads to a possible evolutionary arms race between the MITEs and the host genomes that rapidly and continuously changes the miRNA repertoires. We found that the MITE-miRNAs are selected for targeting genes associated with plant adaptation and habitat expansion, thereby constituting a genomic innovation potentially underlying angiosperm megadiversity.

## Main

MicroRNAs (miRNAs) are an important class of endogenous regulatory RNAs in eukaryotes for programming gene expression. The 20-24 nucleotide (nt) miRNAs arise from immediate precursor transcripts called pre-miRNAs that contain sequences capable of forming the characteristic intra-molecular hairpin structures (Voinnet, 2009; Ma et al., 2010; Yang et al., 2012; Rogers and Chen, 2013; Bartel, 2018). Processed by the evolutionarily conserved double-stranded RNA specific endoribonuclease (Dicer or Dicer-like in plants), the yielded mature miRNAs guide both transcriptional and post-transcriptional gene regulations (Bartel, 2004; Voinnet, 2009; Rogers and Chen, 2013). In plants, numerous studies have firmly established the regulatory function of miRNAs in governing development and stress responses (Voinnet, 2009; Rogers and Chen, 2013; Rodriguez et; al., 2016; Gao et al., 2021). However, the vast plant miRNA repertoires exhibit a striking degree of evolutionary fluidity with only a fraction conserved at different taxonomic levels (Cuperus et al., 2011; Chavez Montes et al., 2014; Guo et al., 2020). Thus, genomic mechanisms for swift *de novo* miRNA origination must be in place, which, when coupled with selection and rapid gene lost, would allow individual lineages to retain largely non-overlapping sets of miRNAs (Chen and Rajewsky, 2007; Moran et al., 2017; Zhang et al., 2021).

In plants, three models of *de novo* miRNA origination have been conceptualized. The target gene inverted duplication model proposes that occasional inverted duplication during gene family expansion might yield transcripts that fold into near-perfect hairpin structures, which are initially processed to produce small interference RNAs (siRNAs). When mutations compromise complementarity of the hairpin structures, they could switch to produce miRNAs (referred to as TID-miRNAs herein) (Allen et al., 2004; Cui et al., 2017; Baldrich et al., 2018). The long terminal repeat (LTR) model predicts that transcribed LTRs of retrotransposons could fold into long hairpin structures triggering production of siRNAs and some might eventually give rise to miRNAs (referred to as LTR-miRNAs) (Piriyapongsa and Jordan, 2008; Voinnet, 2009). The third model is based on miniature inverted-repeat transposable elements (MITEs), which are truncated derivatives of autonomous DNA transposons and contain terminal inverted repeats (TIRs, usually 10-14 base pairs long) flanked by two short direct repeats called target site duplications (TSDs) (Feschotte and Mouchès, 2000; Yang et al., 2001; Feschotte et al., 2003). Their relatively small sizes (generally 50-800 base pairs long) and high copy numbers have led to the proposal that MITEs co-transcribed with the host genes could produce the imperfect hairpin structures for biogenesis of new miRNAs (termed MITE-miRNAs) (Piriyapongsa and Jordan, 2007; Cui et al., 2017). However, these models are only supported by sporadic examples and have not been assessed on a genome scale in different plant species. More important, suitability of these models for continuingly producing novel miRNAs at the expected high evolutionary speed has not been tested.

In this study, by comprehensively integrating information of the miRNA portfolios with genomic annotations, we systemically assessed the three miRNA origination models in 20 phylogenetically representative plant species. The overarching findings illustrated a transposition-transcription process for MITE-miRNA biogenesis and demonstrated MITEs as the predominant genomic source for *de novo* miRNAs in angiosperm. We further found that the rapidly domesticated MITE-miRNAs were selected for targeting genes associated with habitat expansion. These findings add new insights into plant miRNA evolution.

## Results

### Rapid diversification of plant miRNAs

Analyzing the 20,338 miRNAs in 88 plant species phylogenetically ranging from chlorophytes to angiosperms (Guo et al., 2020) revealed 5,022 miRNA families based on homology of the mature miRNAs (Supplementary Fig. 1a). We found that none of the 249 identified algal miRNA families were present in land plants (Supplementary Fig. 1b, c; Supplementary Table 1). Among the 4,773 families in land plants, 4,383 (91.8%) were only found in a single species (Supplementary Fig. 1c). Pairwise comparison of miRNA families among 30 taxonomic families of land plants showed that at least 40% miRNA families were present only in a single family (Supplementary Fig. 1d). In four genera of the Poaceae family, we identified 1,073 miRNA families of which only 27 (2.5%) were present in all four genera while 1,016 (94.7%) were present in a specific genus (Supplementary Fig. 1e; Supplementary Table 2). Within the *Oryza* genus, 202 (41.4%) of the 488 miRNA families in modern *O. sativa* ssp. *japonica* were not present in the other seven examined species or subspecies (Fig. 1a; Supplementary Fig. 2).

**Fig. 1.**
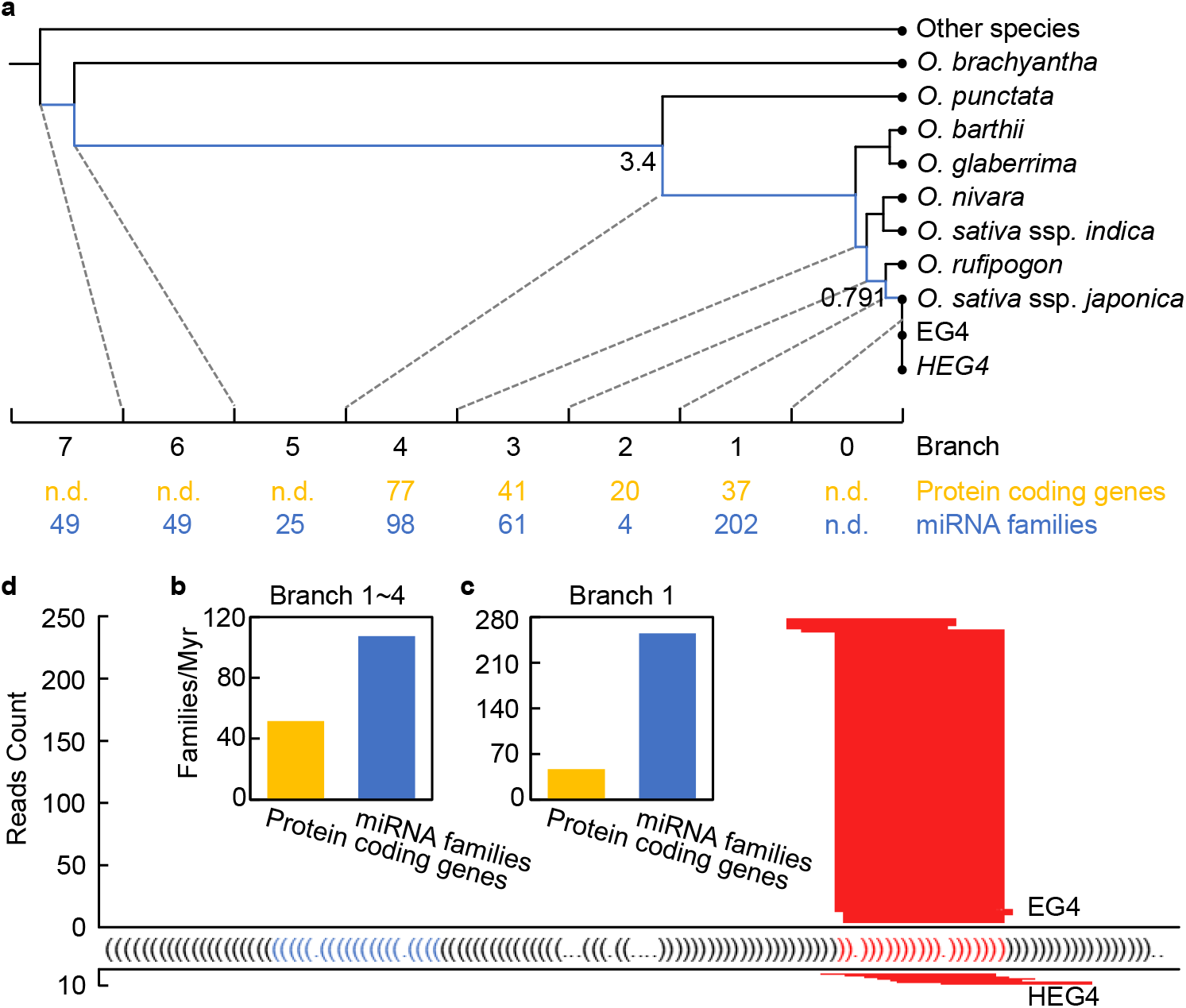
Analysis of de novo miRNA families during *Oryza* diversification. **a,** Evolutionary distribution of origination events for *de novo* miRNA families in the *Oryza* genus towards *O. sativa* ssp. *japonica*. Numbers in blue indicate the number of *de novo* miRNA families identified in each branch. For comparison, distribution of *de novo* protein-coding genes as reported in a previous study (Zhang et al., 2019) is shown in orange. The estimated divergence time was retrieved from the TimeTree database. n.d., not determined. **b-c,** Comparison of the net gain rates of *de novo* protein-coding genes and miRNA families in branch 1 to 4 (**b**) and branch 1 only (**c**). **d,** *Osa-MIRN4438* is unique to EG4. Secondary structure of *Osa-MIRN4438* in EG4 represented with the dot-parenthesis notation in which dots and matching parentheses symbolize base pairs and unpaired bases, respectively. Horizontal red lines represent distribution of the sRNA reads at the *Osa-MIRN4438* genomic locus in EG4 and HEG4.

We traced the rate of net gain of *de novo* miRNA families in the *Oryza* genus over evolutionary time. By examining synteny between closely related *Oryza* genomes and reciprocal-best whole-genome alignments, 175 *de novo* protein-coding genes were identified in *O. sativa* ssp. *japonica* after its divergence from *O. punctata* ∼3.4 million years (Myr) ago (Zhang et al., 2019). By the same comparison, we identified 365 *de novo* miRNA families in *O. sativa* ssp. *japonica*, a net gain of 107.4 new miRNA families per Myr (Fig. 1b), or 2.1 times higher than the net gain of protein-coding genes from ancestral non-coding sequences (Zhang et al., 2019). After the most recent divergence from *O. rufipogon*, the estimated rate of net gain was 255.4 miRNA families per Myr or 5.5 times higher than de novo protein-coding genes (Fig. 1c). To further showcase the swiftness of *de novo* miRNA origination, we compared the miRNA portfolios of EG4 and HEG4, descendants of two single seeds of the *O. sativa* ssp. *japonica* Gimbozu accession EG4* that were maintained as separate lines by self or sibling pollination for about 20 generations (Lu, 2017). *Osa-MIRN4428* with canonical miRNA features was identified in EG4 but not in HEG4 (Fig. 1d), indicating that 20 generations were sufficient to gain a *de novo* miRNA. These observations support previous conclusions of high divergence rates of plant miRNAs (Cuperus et al., 2011; Chavez Montes et al., 2014; Guo et al., 2020), which we further attributed to a rapid origination of novel miRNA families.

### MITE is the dominant source for *de novo* miRNAs in angiosperm

To identify the genomic sources for *de novo* miRNAs, we selected 20 phylogenetically representative species with high-quality reference genomes and available small RNA sequencing (sRNA-Seq) data, including two green algae (*Chlamydomonas reinhardtii* and *Volvox carteri*), one moss (*Physcomitrella patens*), one lycophyte (*Selaginella moellendorffii*), one gymnosperm (*Ginkgo biloba*), one basal angiosperm (*Amborella trichopoda*), five monocots in Poaceae, and nine dicotyledons from five taxonomic families (Supplementary Table 3). In these 20 species, we collected 6,213 high-quality miRNAs from PmiREN1.0 (Guo et al., 2020) and an additional non-redundant set of 1,891 miRNAs from miRBase (Kozomara et al., 2018). Fitting the 8,104 miRNA loci against the three *de novo* origination models using stringent criteria (Supplementary Fig. 3), we identified 341 TID-miRNAs (4.2%) belonging to 248 families, 426 LTR-miRNAs (5.3%) belonging to 168 families, and 1,405 MITE-miRNAs (17.3%) belonging to 557 families (Supplementary Fig. 4a, c; Supplementary Table 4; Supplementary Data 1). The three types of miRNAs were found in both the green alga and land plant lineages (Fig. 2a, b; Supplementary Fig. 5a, b; Supplementary Table 4). Further, most TID-miRNA families (81.0%), LTR-miRNA families (81.0%), and MITE-miRNA families (88.3%) were species specific, indicating that novel miRNAs have heterogeneous originations in plants.

**Fig. 2.**
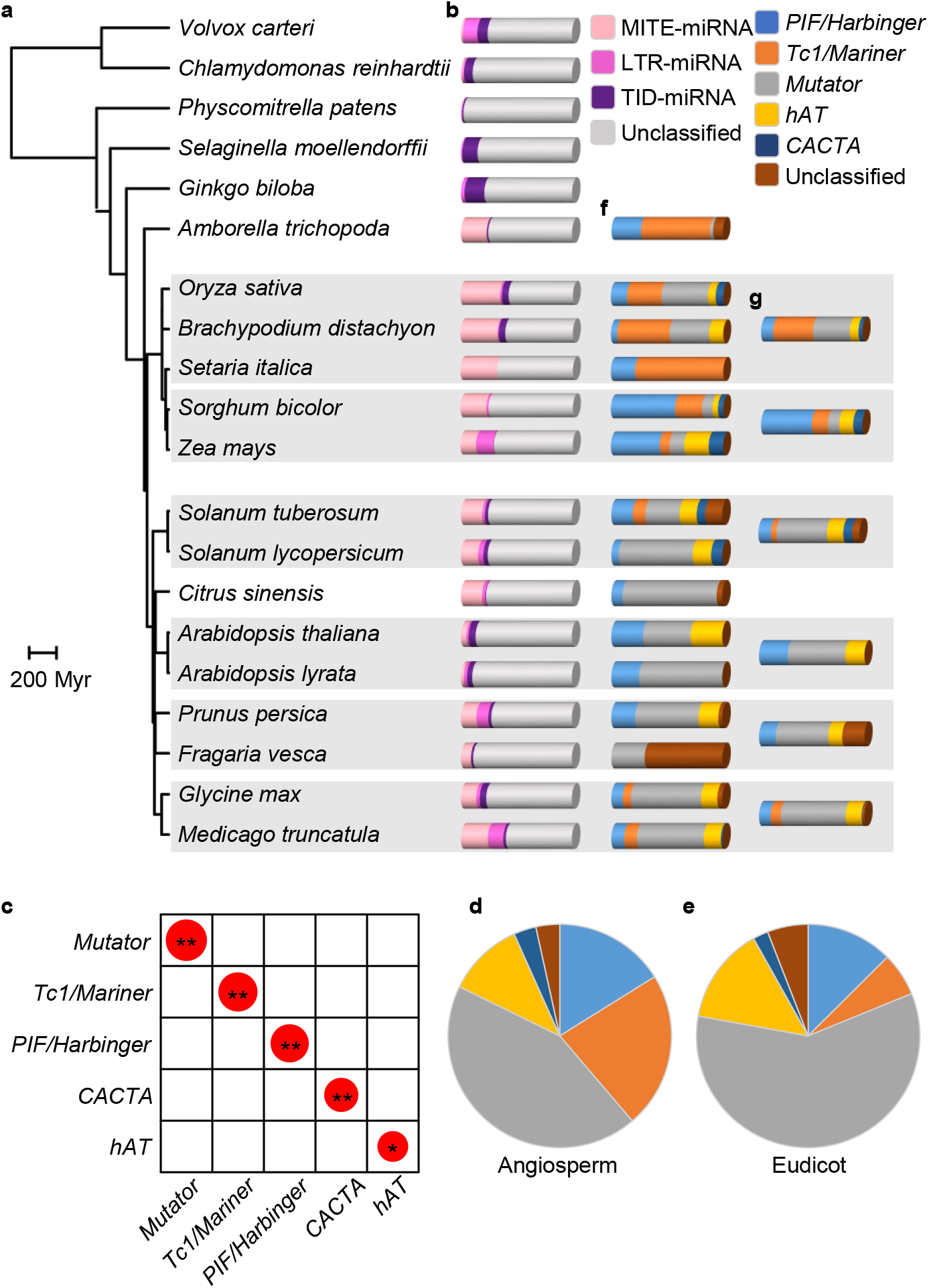
Lineage-specific burst of MITE-miRNAs in angiosperm. **a,** Phylogenetic tree of 20 representative plant species retrieved from TimeTree.org. **b,** Bar graphs showing proportions of MITE-miRNA families, LTR-miRNA families, TID-miRNA families, and unclassified families in each species. **c,** Correlation between the proportion of MITE families belonging to the five indicated superfamilies and proportion of MITE-miRNA families derived from each superfamily in the fifteen examined angiosperm species. Diameter of the circles represents Pearson’s correlation coefficient with the size of the box set to one. *, *P*<0.05, ** *P*<0.01. **d-f,** Proportion of MITE-miRNA families derived from the five MITE superfamilies in all examined angiosperm species (**d**), examined dicots (**e**), and individual species (**f**). **g,** Average proportion of MITE-miRNA families derived from the five MITE superfamilies in the six taxonomic groups as shaded in (**a**).

Among the three types of miRNAs, MITE-miRNAs exhibited the highest numbers, due to a drastic expansion in angiosperms (Fig. 2a, b). Only six, or less than 0.5% miRNAs were MITE-miRNAs in the five examined non-angiosperm plants: three in *V. carteri*, two in *C. reinhardtii*, one in *P. patens* (Fig. 2b; Supplementary Fig. 5b; Supplementary Table 4). In contrast, 1,399 (20.6%) miRNAs, or 552 (16.9%) miRNA families, were identified as MITE-miRNAs in the 15 examined angiosperms (Supplementary Fig. 4b, d), or a burst of two orders of magnitude over non-angiosperms. In the basal angiosperm *A. trichopoda*, 59 or 25.0% miRNAs (34 or 23.3% families) were MITE-miRNAs, indicating that the burst of MITE-miRNAs had already occurred in early angiosperms (Fig. 2b; Supplementary Fig. 5b; Supplementary Table 4).

Using the *Oryza* genus as an example, we compared the rates of net gain of the three types of miRNAs. In *O. sativa*, we identified 44 TID-miRNA, 10 LTR-miRNA and 118 MITE-miRNA families (Fig. 2b; Supplementary Fig. 6). We found that the rates of net gain of MITE-miRNA families were consistently higher than LTR-miRNA and TID-miRNA families throughout evolution of the *Oryza* genus (Supplementary Fig. 6). Further, we confirmed that *Osa-MIRN4428*, the *de novo* miRNA in EG4 after its split from HEG4 (Fig. 1d), was a MITE-miRNA (Supplementary Fig. 7). *Osa-MIRN4428* is homologous to 37 copies of an active MITE family belonging to the *Mutator* superfamily (Supplementary Fig. 7). Sequence alignment revealed that *Osa-MIRN4428* accumulated five single nucleotide polymorphisms (SNPs) compared to the related MITEs (Supplementary Fig. 8). However, only the SNP in the mature miRNA region of *Osa-MIRN4428* created a mismatch in the secondary structure (Supplementary Fig. 8), suggesting that this SNP was responsible for converting the MITE to a miRNA. Collectively, these observations demonstrate that MITEs are the dominant source for rapid *de novo* miRNA origination in *Oryza*, a conclusion corroborated by analyzing the *Solanum* genus (Supplementary Fig. 6).

It should be noted that the above-reported MITE-miRNAs were identified based on high stringent criteria requiring extensive sequence homology between MITE-miRNAs and the founding MITEs. As homology would be eroded over evolutionary time, some MITE-miRNAs may have escaped detection by current methods (Moran et al., 2017). In fact, lowering the requirement of the minimal matched length substantially increased the number of putative MITE-miRNAs. For example, we found evidence supporting a MITE origination of miR156, one of seven miRNA families deeply conserved in land plants (Cuperus et al., 2011; Chavez Montes et al., 2014; Guo et al., 2020). In both *O. sativa* and the model dicot *Arabidopsis thaliana*, we identified a member of the miR156 family (*Osa-MIR156G* and *Ath-MIR156D*) that shares similarity with MITEs in the *Mutator* superfamily (Supplementary Fig. 9). Synteny analysis revealed that both loci were of recent origins. For example, the *Osa-MIR156G*-containing fragment emerged in both *O. sativa* and *O. rufipogon* about 0.791 Myr ago (Supplementary Fig. 9). Thus, MITE is the potential source for even the most conserved miRNA families in land plants.

### Diversification of MITE families influences frequency of MITE-miRNAs

Within the examined angiosperm species, no correlation was found between the number of MITE-miRNA families (Supplementary Table 4) and the number of MITEs (Supplementary Table 5) (Pearson’s r = 0.005, *P* = 0.767). This observation prompted us to examine the MITE families. Based on homology of the TIR and TSD, MITEs in the green plant lineage are divided into seven superfamilies, including *Tc1/Mariner*, *PIF/Harbinger*, *Mutator*, *CACTA*, *hat*, *P element* and *Novosib* (Feschotte and Mouchès, 2000; Fattash et al., 2013; Chen et al., 2014). The examined plants exhibited drastically different composition of the seven superfamilies (Supplementary Fig. 10; Supplementary Table 5, 6). Consistent with previous reports (Chen et al., 2014), *P element* and *Novosib* were only found in *C. reinhardtii* and thus had little impact on the MITE-miRNAs. In the other five superfamilies, the proportion of MITEs and the proportion of MITE-miRNA families derived from each superfamily showed a strong correlation (Fig. 2c; Supplementary Table 7).

We found that the *hAT* superfamily was dominant in the five non-angiosperm species in which *Mutator* and *CATCA* were absent. In contrast, *Mutator* was the dominant superfamily in dicots while *PIF/Harbinger* and *Tc1/Mariner* were dominant in the basal angiosperm and monocots (Supplementary Fig. 10; Supplementary Table 5). Consequently, we found that an overwhelming portion (81.7%) of MITE-miRNA families in angiosperm were derived from *Mutator* (43.1%), *Tc1/Mariner* (22.4%), and *PIF/Harbinger* (16.2%) (Fig. 2d; Supplementary Table 8, 9). Furthermore, over half MITE-miRNA families (59.0%) in dicots were derived from *Mutator* alone (Fig. 2e). In the examined monocots, 39.4% MITE-miRNA families in the clade consisting of *O. sativa*, *Brachypodium distachyon*, and *Setaria italica* were derived from *Tc1/Mariner* whilst 50% MITE-miRNA families in the clade containing *Sorghum bicolor* and *Zea mays* were from *PIF/Harbinger* (Fig. 2f, g).

Within each superfamily, MITEs of sequence homology were further grouped into families. After annotating the MITE families, we found that the number of MITE families in angiosperm was generally an order of magnitude higher than that in non-angiosperm (Supplementary Table 6). Comparing the number of MITE families and MITE-miRNA families revealed a significant positive linear correlation (Pearson’s r = 0.546, *P* < 0.01) (Supplementary Fig. 11). We found that 70.7% MITE families in angiosperms belonged to *Mutator*, *Tc1/Mariner* and *PIF/Harbinger*, while the proportion was 35.7% in non-angiosperms (Supplementary Table 6). These results indicate that the drastic increase in MITE-miRNAs in angiosperms correlates with expansion of the *Mutator*, *Tc1/Mariner* and *PIF/Harbinger* superfamilies.

### Intrinsic MITE features determine conversion to miRNAs

To identify the features of *Mutator*, *Tc1/Mariner*, and *PIF/Harbinger* that impact their conversion to miRNAs, we tested four parameters, including copy number of family members (which reflects their transposition activity), normalized minimal free energy (NMFE) of the MITE transcripts (which reflects stability of the hairpin structures), the guanine-cytosine (GC) content, and length of the MITE loci. We divided the *Mutator*, *Tc1/Mariner*, and *PIF/Harbinger* MITEs into two groups, the MITE-miRNA matching MITEs (mMITEs) and MITEs that did not match the MITE-miRNAs (nMITEs). We found that the mMITEs and nMITEs exhibited significant difference regarding the four parameters (Supplementary Fig. 12a-d). Using a machine learning approach based on a gradient boosting decision tree algorithm (Chen and Guestrin, 2016), we assessed relative importance of these parameters on the conversion to MITE-miRNAs. While this analysis achieved an overall accuracy of 96.8%, copy number of MITE families was the most influential feature (Supplementary Fig. 12e). Further analysis revealed that proportion of the mMITEs with over 400 members was significantly higher than that of the nMITEs (Supplementary Fig. 12f), suggesting that transposition activity of MITEs is pivotal for MITE-miRNA formation.

High importance scores were also retrieved for length and NMFE of MITEs (Supplementary Fig. 12e), suggesting that the ability to form hairpin RNA structure is important for the generation of MITE-miRNAs. Further comparison revealed that length of MITEs has a significant impact on the miRNA portfolios of closely related species. *Z. mays* and *S. bicolor* of Poaceae diverged from a common ancestor about 12.18 Myr ago. Although MITEs in *Z. mays* (764 copies on average) were more active than those in *S. bicolor* (408 copies on average), number of MITE-miRNA families was quite close in *Z. mays* (21) and in *S. bicolor* (23). While NMFE of the MITEs (Supplementary Fig. 13) and length distribution of MITE-miRNAs (Fig. 3a) were comparable in the two species, length of MITEs in *Z. mays* (maximum of 132 bp) was drastically shorter than that in *S. bicolor* (maximum of 244 bp) (Fig. 3a). Reduction in MITE length occurred in both *PIF/Harbinger* (Fig. 3b) and *Tc1/Mariner* (Fig. 3c), the two dominant superfamilies in *Z. mays* and *S. bicolor*. Further analysis revealed that the length reduction was caused by a burst of short MITEs. In *S. bicolor*, only 15 MITEs in *PIF/Harbinger* and seven MITEs in *Tc1/Mariner* were shorter than 100 bp. By contrast, the numbers jumped to 7,663 in *Z*. *mays*, or 4.2% of *PIF/Harbinger* and 18.5% of *Tc1/Mariner*. Thus, burst of extremely short MITEs, which likely imposes constraints on forming productive RNA hairpin structures, impeded the origination of MITE-miRNAs in *Z*. *mays*.

**Fig. 3.**
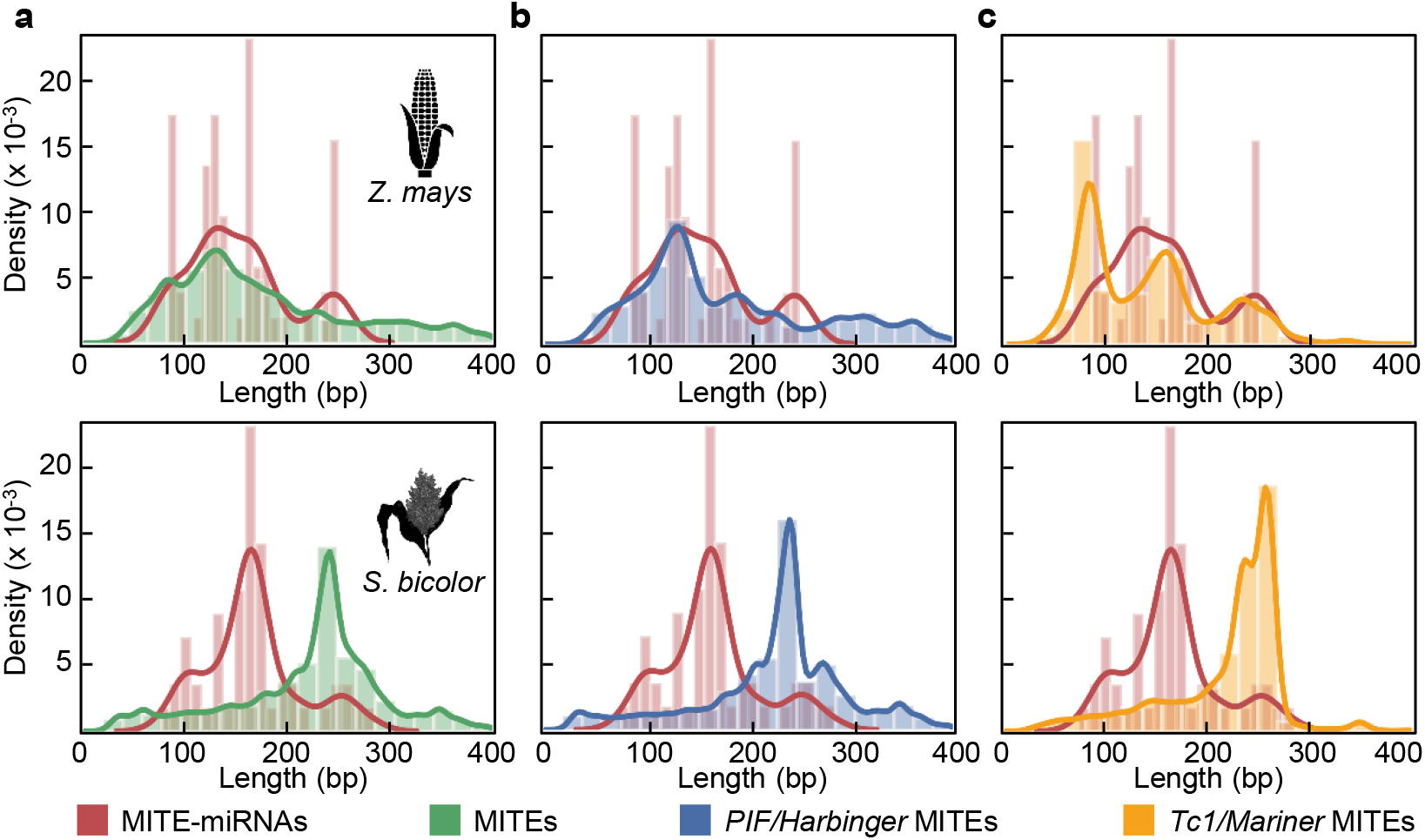
Length distribution of MITE-miRNAs and MITEs in *Z. mays* and *S. bicolor*. **a,** Comparison of the length distribution of MITE-miRNAs and MITEs in *Z. mays* (top) and *S. bicolor* (bottom). Kernel density estimation with histograms is used to display the length distribution. **b,** Comparison of the length distribution of MITE-miRNAs and MITEs from the *PIF/Harbinger* superfamily in *Z. mays* and *S. bicolor*. **c,** Comparison of the length distribution of MITE-miRNAs and MITEs from the *Tc1/Mariner* superfamily in *Z. mays* and *S. bicolor*.

A similar pattern of length distribution was observed for the dominant *Mutator* superfamily in two Fabaceae species, *Medicago truncatula* and *Glycine max* (Supplementary Fig. 14). While MITEs in *G. max* (1,344 copies on average) was more active than those in *M. truncatula* (461 copies on average), *G. max* (68) contained less MITE-miRNA families in comparison to *M. truncatula* (78). Again, no significant difference in NMFE was found (Supplementary Fig. 13), but the length of *Mutator* MITEs was drastically longer in *M. truncatula* (maximum of 183 bp) than in *G. max* (maximum of 92 bp) (Supplementary Fig. 14). In conclusion, within *Mutator*, *Tc1/Mariner*, and *PIF/Harbinger* MITEs, copy number and length are two critical parameters determining the origination of MITE-miRNAs.

### A transposition-transcription process for MITE-miRNA biogenesis

To gain insight into the process through which MITE-miRNAs are generated, we analyzed genomic distribution of MITE-miRNAs in the 15 examined angiosperms. We divided the genomes into four compartments based on current annotation, including exon, intron, promoter, and intergenic region. We found that the proportion of MITE-miRNAs in introns was significantly higher than non-MITE-miRNAs whereas proportion of MITE-miRNAs in exons was significantly lower than non-MITE-miRNAs (Fig. 4a). Calculation of the length-normalized proportions of MITE-miRNA and other miRNAs in different genomic compartments confirmed that MITE-miRNAs were preferentially located in introns (Fig. 4b), particularly the first introns (Supplementary Fig. 15). Analysis of RNA-sequencing data in *O. sativa* revealed that the transcript levels of the MITE-miRNA host genes (in which a MITE-miRNA locates), especially those hosting the MITE-miRNAs in introns, were significantly higher than host genes of nMITEs (Supplementary Fig. 16). These findings indicate that MITE-miRNAs are preferentially located in the first intron of actively transcribed host genes.

**Fig. 4.**
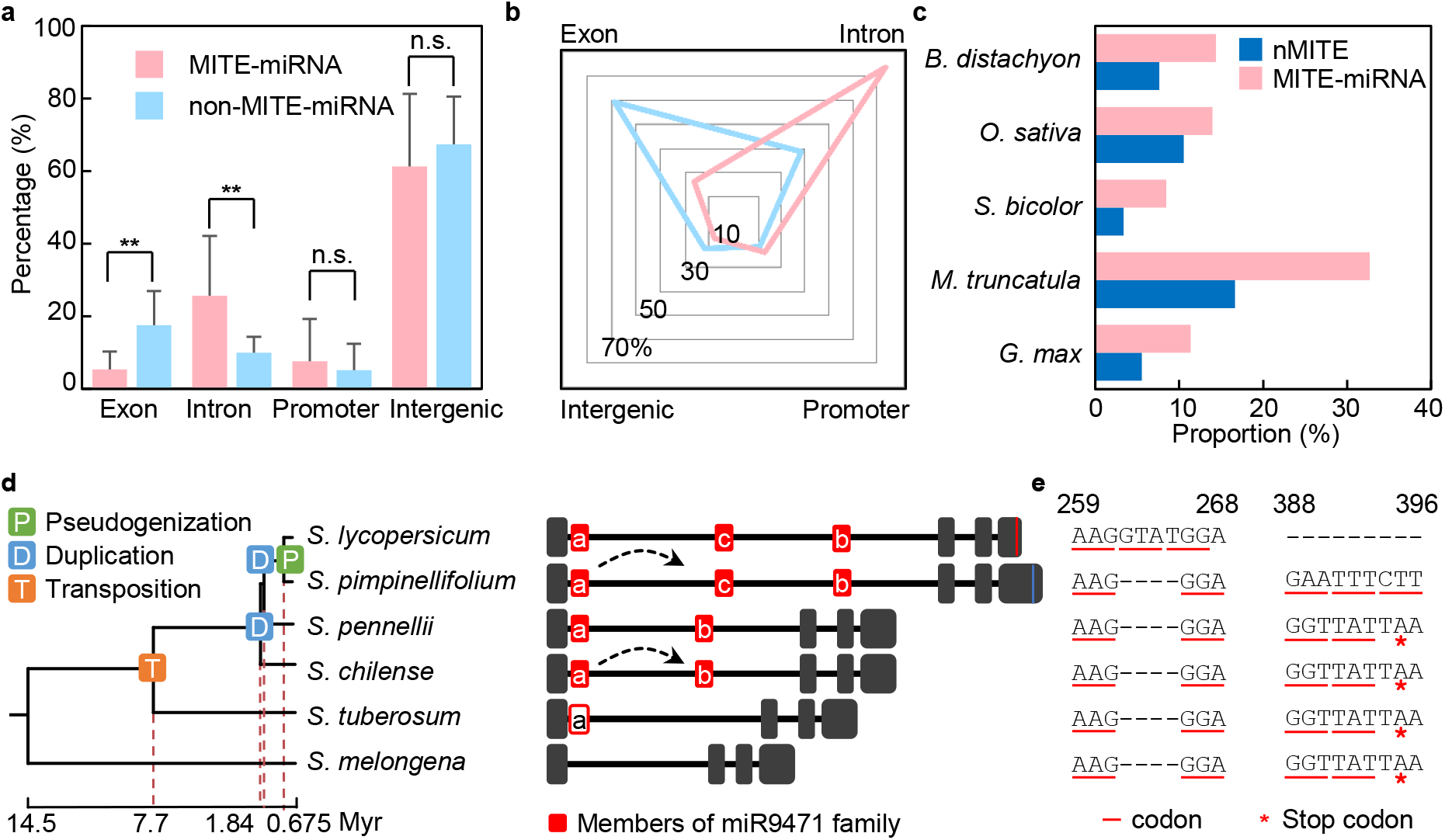
Origination of MITE-miRNAs through a transposition-transcription process. **a,** Histogram showing the proportion of MITE-miRNAs located in exons, introns, promoters, and intergenic regions. **, *P* < 0.01 by independent samples t-test. n.s., not significant. **b,** Length normalized genomic distribution of MITE-miRNAs and other miRNAs in exons, introns, promoters, and intergenic regions. **c,** Proportion of MITE-miRNAs and nMITEs associated with annotated pseudogenes in the five examined species. **d,** Evolutionary trajectory of *MIR9471* and its host gene in *Solanum*. The phylogenetic tree and divergence time were retrieved from TimeTree.org. Structures of the host genes corresponding to the given species are shown right to the tree. Exons are represented by black boxes and introns by horizontal lines. Solid red boxes mark positions of the miR9471 family members in the first intron of the host genes. Open red box indicates a sequence in *S*. *tuberosum* homologous to *MIR9471* but lacks evidence as a miRNA. Dotted arrows represent two tandem duplications that expanded the miR9471 family. Vertical lines in the last exons of *S. lycopersicum* and *S. pimpinellifolium* indicate mutations that pseudogenized the host gene. Order of these evolutionary events was deduced by sequence comparison among the species and indicated on the tree with colored squares. **e,** Sequence alignments at two sites in the host gene of *MIR9471* where the four-nucleotide-insertion occurs in *S. lycopersicum* and mutations disrupting the stop codon occur in *S. pimpinellifolium*, respectively. Numbers at the top indicate positions in the host gene transcript.

A MITE-miRNA nested inside a host gene would create a conflict as the two functional entities may require different spatiotemporal expression. We hypothesized that one way to resolve the conflict is host gene pseudogenization and thus examined five species with comprehensive annotation of pseudogenes, including three grass species *B. distachyon*, *O. sativa*, and *S. bicolor*, and two legume species *M. truncatula* and *G. max* (Xie et al., 2019). We found that MITE-miRNAs were indeed more frequently overlapped with pseudogenes than nMITEs (Wilcoxon signed-rank test, *P* < 0.5) (Fig. 4c). The miR9471 family in the genus *Solanum* afforded an example for illustrating the process of MITE-miRNA associated pseudogenization. Reconstruction the evolutionary history of miR9471 revealed an initial transposition of a MITE (SQ382168580 of the *Tc1/Mariner* superfamily) into the first intron of *Solyc12g008590* (encoding the highly conserved profilin) in *Solanum tuberosum* after its divergence from *S. melongena* 14.5∼7.7 Myr ago (Fig. 4d; Supplementary Fig. 17). The family was expanded by two tandem duplication events occurring 7.7∼1.84 Myr ago in *S. chilense* and *S. pennellii* and 1.80∼0.675 Myr ago in *S. pimpinellifolium* and *S. lycopersicum*, respectively (Fig. 4d). Congregation of miR9471 members in the first intron of *Solyc12g008590* was concurrent with a specific expression of miR9471 at the green-to-breaker stage transition during fruit development while the host gene exhibits constitutive expression (Supplementary Fig. 18). This event was followed by convergent pseudogenization of *Solyc12g008590*, occurred in *S. lycopersicum* as a frame shift caused by an insertion of four base pairs in the fourth exon that led to a premature stop codon, and in *S. pimpinellifolium* as missense mutations that abolished the original stop codon (Fig. 4e; Supplementary Fig. 19). These findings indicate that selection of the MITE-miRNAs may have a collateral selection of the host gene as a non-coding miRNA precursor, which would lead to pseudogenization to resolve conflict with the original function of the host gene.

### Modern MITE-miRNAs facilitate angiosperm habitat expansion

In the 20 examined plant species, we identified 58,038 putative target genes for the 8,095 miRNAs (Supplementary Data 2). Consistent with previous reports that TID-miRNAs are derived from and target large gene families (Baldrich et al., 2018), we observed that the average number of target genes per TID-miRNA (10.4) was significantly higher than that of MITE-miRNA (7.5) and LTR-miRNA (5.5) (Supplementary Fig. 20). Gene Ontology (GO) analysis revealed that molecular functions of the MITE-miRNA target genes, compared to those of the TID-miRNAs and LTR-miRNAs, were more enriched with terms related to responses to environmental stimuli (Fig. 5a; Supplementary Fig. 21-23). Among the most conspicuous environmental factors associated with the MITE-miRNA target genes was temperature, as the GO terms “response to temperature stimulus”, “response to freezing”, “response to cold”, and “response to heat” were all significantly enriched (Supplementary Fig. 24).

**Fig. 5.**
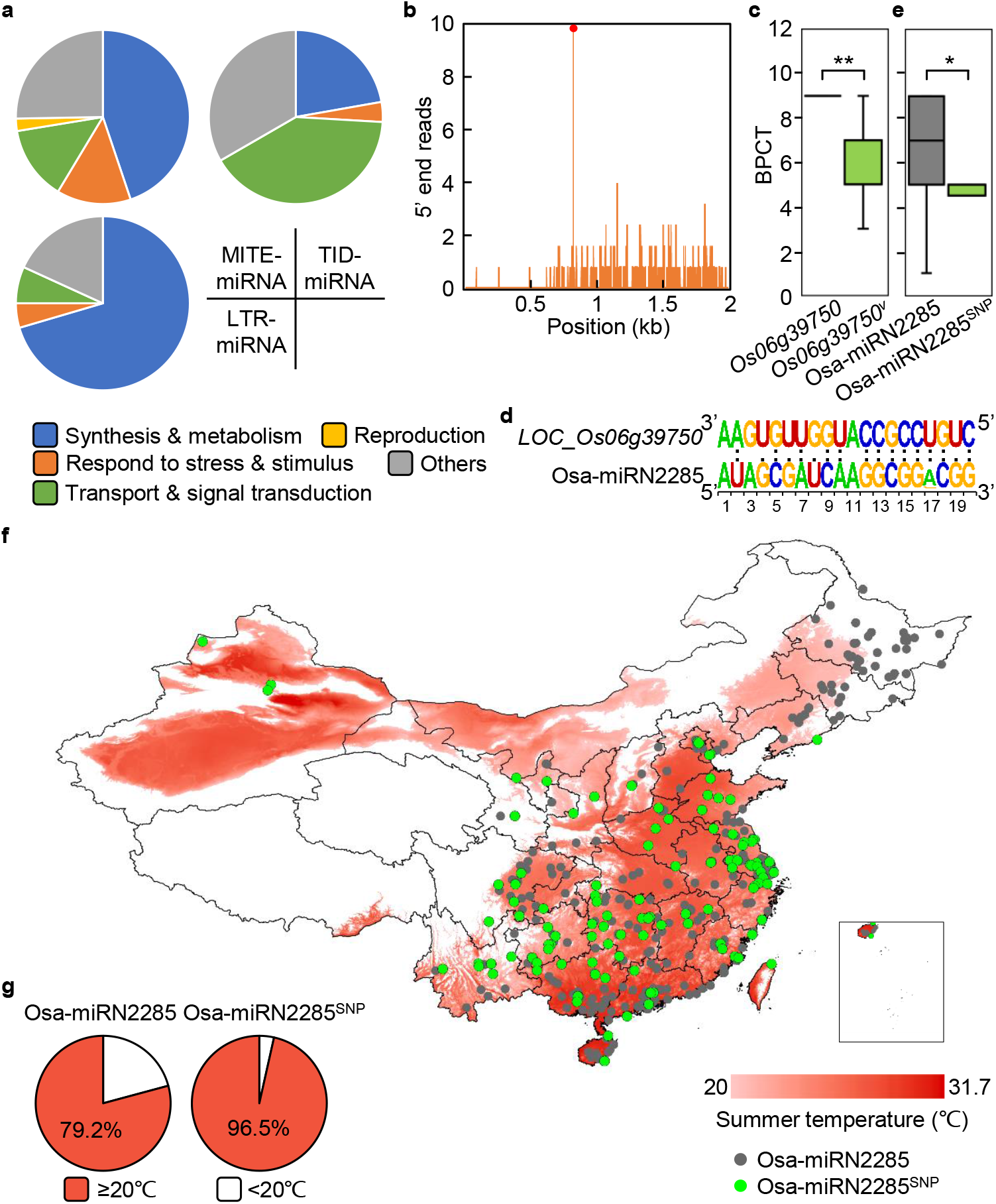
MITE-miRNAs facilitate habitat expansion of the angiosperm. **a,** Pie charts showing the proportions of enriched GO terms associated with target genes of the MITE-miRNAs, LTR-miRNAs, and TID-miRNAs. **b,** Validation of *LOC_Os06g39750* as an authentic Osa-miR2285 target by degradome sequencing. The horizontal and vertical axes represent the position of the *Os06g39750* transcript and the frequency of sequenced transcript ends, respectively. Red dot indicates the identified Osa-miRN2285-guided cleavage site. **c,** Comparison of BPCT between the *Os06g39750* type (n = 14) and *Os06g39750^v^* type (n = 48) *O*. *sativa* germplasms. **, *P* < 0.01 by independent samples t-test. **d,** Weblogo showing base complementarity between Osa-miRN2285 and its target site in *Os06g39750*. Data were compiled from sequences in 5,459 germplasms. **e,** Comparison of BPCT between the Osa-miRN2285 type (n = 76) and Osa-miRN2285^SNP^ type (n = 6) germplasms. *, *P* < 0.05 by independent samples t-test. **f,** Comparison of the geographical distribution of *O*. *sativa* germplasms with summer temperature in China. Geographical origins of the Osa-miRN2285 type (grey) and Osa-miRN2285^SNP^ type (green) germplasms are marked on the map. Distribution of the ten-year average summer temperature is superimposed on the map with temperature below 20°C shown in white and temperature above 20°C in different shades of red. **g,** Proportions of the Osa-miRN2285 type and Osa-miRN2285^SNP^ type germplasms produced in areas with high (≥20°C) and low summer temperature (<20°C).

We selected Osa-miRN2285 in *O*. *sativa*, which targets *LOC_Os06g39750* based on degradome sequencing data (Fig. 5b), for analyzing the role of a MITE-miRNA in temperature responses. We found three lines of evidence supporting the involvement of the Osa-miRN2285-*Os06g39750* circuit in cold adaption. First, *Os06g39750* was known to be induced by cold-water treatment (Sun et al., 2018). Expression of Osa-miRN2285 exhibited the reciprocal pattern and significantly decreased after cold treatment (Supplementary Fig. 25). Second, *Os06g39750* was a candidate gene for cold tolerance based on a quantitative trait loci analysis (Sun et al., 2018). By analyzing re-sequencing data and phenotypic records from public database (Peng et al., 2019), we found that *O*. *sativa* germplasms containing sequence variations in *Os06g39750* (the *Os06g39750^v^* type) exhibited significantly lower (*P* < 0.01 by independent samples t-test) Budding Phase Cold Tolerance (BPCT) than germplasms with reference *Os06g39750* sequence (the *Os06g39750* type) (Fig. 5c). Finally, we detected a SNP at the 17^th^ position in mature Osa-miRN2285 (Fig. 5d). Out of 5,459 examined germplasms, 993 possessed this SNP undermining binding to *Os06g39750* and were referred to as the miRN2285^SNP^ type. Phenotypic records showed that the miRN2285^SNP^ type germplasms exhibited BPCT significantly lower (*P* < 0.05 by independent samples t-test) than the miRN2285 type germplasms (Fig. 5e).

To demonstrate that miRN2285 facilitates adaptation to colder habitat, we collected geographical information of the production sites of different *O*. *sativa* germplasms in China and calculated the ten-year (from 2009 to 2018) mean summer temperature. Superimposing the geographical and temperature distribution showed that production sites of the miRN2285 type germplasms included the northeast region that has lower summer temperature (Fig. 5f). Moreover, the local summer temperature at the production sites of the 171 miRN2285^SNP^ type germplasms was significantly higher than that of the 1,177 miRN2285 type germplasms (Fig. 5g; Supplementary Fig. 26). These results suggest that miRN2285 is selected in regions with lower growing season temperature and the selection is relaxed when these germplasms are introduced to regions with higher temperature. Collectively, our findings suggest that MITE-miRNAs are pertinent to habitat expansion of the angiosperms by selectively target genes related to environmental adaption.

## Discussion

Biogenesis of miRNAs from a DNA segment requires its ability to be transcribed and sequences that could fold into the hairpin secondary structure (Cui et al., 2017; Baldrich et al., 2018). Our findings provided genome-scale support for miRNA origination from MITEs that meets these requirements (Fig. 6). MITEs transpose through the action of transposases encoded by autonomous DNA transposons (Jiang et al., 2003; Yang et al., 2009) and are often found close to or within genes (Kuang et al., 2009; Ohmori et al., 2008; Santiago et al., 2002), allowing TIR-seeded transcripts to fold into imperfect hairpin structures typical of miRNA precursors (Cui et al., 2017). We presented four lines of evidence showing that this process was highly efficient to generate novel miRNAs in angiosperm. First, we found as few as 20 generations and as few as one SNP could be sufficient to allow a MITE-miRNA to arise from a MITE under certain circumstances (Fig. 1d; Supplementary Fig. 7, 8). We also found evidence suggesting some of the deeply conserved miRNA families during plant evolution (e.g., miR156) may have arisen from MITEs (Supplementary Fig. 9). Next, we showed that MITE-miRNA families accumulated to quantities more than the TID-miRNA and LTR-miRNA families combined (Fig. 2; Supplementary Fig. 4, 5). Finally, we found that generation of MITE-miRNAs was often associated with host gene pseudogenization (Fig. 4d, e; Supplementary Fig. 17-19). These findings indicate that MITE became the dominant source for *de novo* miRNA origination in angiosperms through a transposition-transcription process and profoundly shaped the modern miRNA repertoires.

**Fig. 6.**
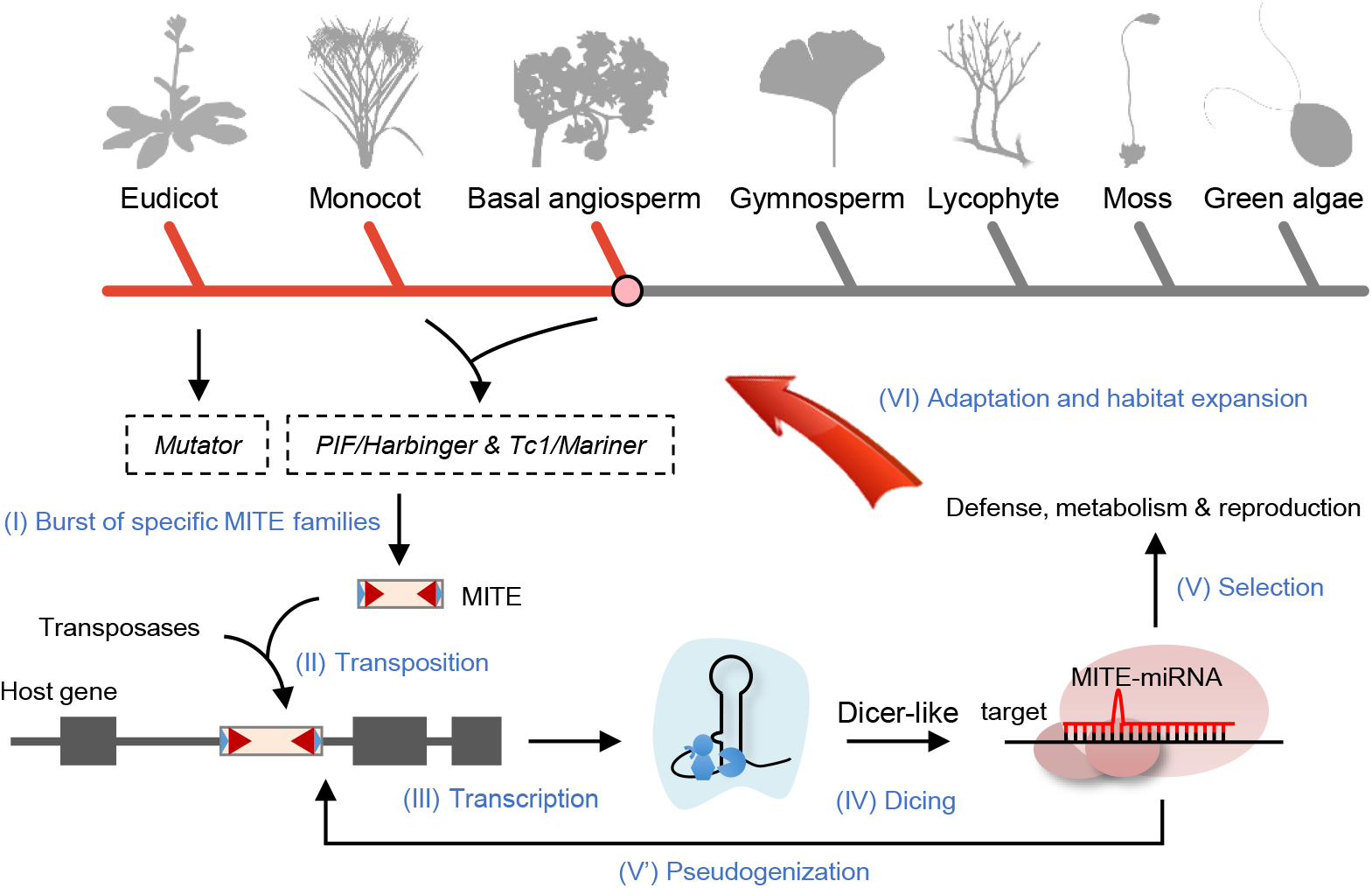
A model for MITEs in shaping the miRNA repertoires in angiosperm. MITEs predispose the miRNA portfolio in angiosperm to diversify through processes I to VI. Bursts of the *Mutator*, *Tc1/Mariner*, and *PIF/Harbinger* MITE families in angiosperms (I) increase the frequency of appropriately sized MITEs to transpose into introns (II). These MITEs hitchhike host gene transcription (III), allowing TIR-seeded regions to fold the transcripts into hairpin structures for the Dicer-like complex to generate *de novo* miRNAs (IV). Selection of the miRNAs through their target genes (V) might have a collateral selection of the host gene as a non-coding miRNA precursor through pseudogenization (V’). Consequently, novel MITE-miRNAs preferentially regulate target genes with biological functions related to habitat expansion and adaptation (VI).

Transposable elements are selfish genetic elements imposing intra-genomic conflicts that can lead to an evolutionary arms race with the host genomes (Werren, 2011; Luo et al., 2020). Our findings indicate that the non-autonomous MITEs could be molecularly domesticated by the evolutionarily conserved Dicer-like protein machinery to produce miRNAs. We found positive correlations between the number of MITE-miRNA families and the diversity of MITE families (Supplementary Fig. 11), which suggests that lineage-specific bursts of MITE-miRNAs in angiosperms was due to the increasing diversity of the MITE families. A model constructed by machine learning revealed that both copy number and length of the MITEs have major effects on the origination of MITE-miRNAs (Supplementary Fig. 12). In addition, we found that MITEs transposed into the first intron of the host genes were mostly likely to become miRNAs (Fig. 4; Supplementary Fig. 15, 16). These findings indicate that conversion to miRNAs was associated with intrinsic features of the MITEs. Consequently, this domestication process serves the host genome by repressing the newly invaded MITEs as well as expanding the miRNA repertoires to enhance gene regulation capacity during evolution (Fig. 6). On the other hand, MITEs may have developed a counter strategy to evade the miRNA-oriented domestication. For instance, bursts of extremely short MITEs may undermine the success of MITE domestication and reduce miRNA origination (Fig. 3; Supplementary Fig. 14). This finding suggests an evolutionary arms race between expansion of the MITEs and the host genome’s efforts of turning them into domestic gene regulators, which provides an explanation why the miRNA repertoire is continuously changing in plants (Fig. 1; Supplementary Fig. 6). Further analysis of the genomic processes associated to *de novo* origination of miRNAs should provide more insights into the nature of genomic conflicts.

Since emergence on land, angiosperms have exceptionally radiated and surpassed all other plant lineages in both species richness and ecological dominance (Rubinstein et al., 2010; Magallón et al., 2013). Currently, angiosperms constitute the dominant vegetation on the Earth’s terrestrial surface and are indispensable to agriculture. Many genetic and genomic studies have attempted to identify the drivers of angiosperm megadiversity, which is thought to involve multivariate interactions among intrinsic traits and extrinsic forces (Vamosi and Vamosi, 2011; Hausch et al., 2018; Magallón et al., 2018; Sauquet and Magallón, 2018; Vamosi et al., 2018). Our findings of massive domestication of MITEs as MITE-miRNAs (Fig. 6), which are selected for targeting genes associated with environmental adaptation and habitat expansion (Fig. 5; Supplementary Fig. 21-26), suggest that MITE-miRNAs have the potential for driving angiosperm diversification. Continuingly and quickly emerging MITE-miRNAs, once incorporated into the gene regulatory network in a lineage-specific manner (Chen and Rajewsky, 2007; Moran et al., 2017; Zhang et al., 2021), would spur rapid buildup of genetic diversity that allows novel traits to evolve over the long term. Therefore, MITE-miRNAs warrant further studies as candidates for genomic innovations underlying angiosperm megadiversity.

## Methods

### Analysis of miRNA conservation and divergence

Sequences and annotations of 20,388 miRNAs in 88 species were retrieved from PmiREN1.0 (https://www.pmiren.com) (Guo et al., 2020). A built-in Perl script, merge_and_rank.bash, in miRDeep-P2 (Kuang et al., 2018) was used to group the miRNA families by allowing no more than two mismatches between the members. Conservation of a miRNA family in each taxon was declared if the miRNA was detected in at least half of the examined species in that taxon. Species-specific miRNA families were defined as those were detected in only one species. The 30 taxonomic families of land plants used for pairwise comparison of miRNA families included Actinidiaceae, Amborellaceae, Araceae, Asparagaceae, Asteraceae, Brassicaceae, Bryophytes, Caricaceae, Caryophyllaceae, Convolvulaceae, Cucurbitaceae, Euphorbiaceae, Fabaceae, Fagaceae, Lythraceae, Lauraceae, Malvaceae, Musaceae, Nelumbonaceae, Oleaceae, Phrymaceae, Pinaceae, Poaceae, Rosaceae, Rutaceae, Salicaceae, Selaginellaceae, Solanaceae, Vitaceae, Zosteraceae. This alphabetic order was followed from top to bottom and from left to right in Supplementary Fig. 1d. Multiple in-house Perl scripts were used to compare the miRNA families among four genera and different species.

### Identification of new miRNA origination events in *Oryza* and *Solanum*

Following the rationale showing in Supplementary Fig. 2, the 428 miRNA families in *O. sativa* ssp. *japonica* were compared to those in 84 species outside of *Oryza*. The 379 miRNA families identified as *O. sativa* specific were further compared to the miRNA portfolios of seven closely related *Oryza* species using sRNA-Seq data. Briefly, 59 sRNA-Seq datasets (Supplementary Table 10) were parsed using Trim Galore (version 0.5.0) (http://www.bioinformatics.babraham.ac.uk/projects/trim_galore/) with parameters ‘-length 18 -max_length 28 -small_rna -stringency 3’ to remove adapters and Bowtie (1.1.2) (Langmead, 2010) with parameters ‘-v 0’ to map clean reads corresponding to mature miRNAs. The ancestral states of the miRNA families were assigned following the parsimony rule previously described for identifying evolutionary events for *de novo* protein-coding genes (Zhang et al., 2019). The generation time of a given miRNA family was determined by the divergence time of species in which the miRNA first appeared. The same method was employed to analyze the origination events of novel miRNA families in *Solanum* using 54 sRNA-Seq datasets from six species (Supplementary Table 11).

### Profiling miRNAs in EG4 and HEG4

sRNA-Seq datasets of EG4 (SAMN03135602; SRR5573658, SRR5573660, SRR5573661) and HEG4 (SAMN03140295; SRR5573662, SRR5573663, SRR5573670) were downloaded from NCBI Sequence Read Archive (SRA). SRA toolkit (version 2.8.2) (Kodama et al., 2011) was used to convert the original compressed files into Fastq format. Trim Galore (version 0.5.0) was used to further trim adapter sequences. Identical reads were collapsed using previously described methods (Kuang et al., 2018; Yang and Li, 2011). Newly updated plant miRNA criteria (Axtell and Meyers, 2018) were employed to identify miRNA candidates based on the *O*. *sativa* ssp. *japonica* reference genome (MSU7 release 7.0). All miRNA candidates were further annotated as previously described (Guo et al., 2020) and miRNAs not registered in miRBase (Kozomara et al., 2018) were marked with the “miRN” prefix.

### Phylogenetic tree and divergence time

The phylogenetic tree of *Oryza* and *Solanum* were retrieved from the TimeTree database (http://timetree.org/) (Kumar et al., 2017; Zhang et al., 2019). TimeTree was also used to estimate the species divergence time in the two genera.

### Identification of MITE-miRNAs

Genome assemblies and annotations of the 20 examined plants are summarized in Supplementary Table 3. For these species, sequences and annotation of 6,213 miRNAs were obtained from PmiREN1.0 (Guo et al., 2020). The MITE sequences in 18 species were directly retrieved from the P-MITE Database (http://pmite.hzau.edu.cn/) (Chen et al., 2014) (Supplementary Table 5, 6). MITEs in *A. trichopoda* and *G. biloba* were *de novo* identified from the genomes using MITE-hunter (Han and Wessler, 2010). The outputs of MITE-hunter were manually checked for TIR and TSD sequences. Only groups with precise boundaries and TIR sequences were further considered. The consensus sequences of homologous MITEs were grouped into families and used as references to scan the entire genomes to find more MITEs using RepeatMasker (http://www.repeatmasker.org/) with the parameter ‘-x -no_is –nolow -cutoff 250’. The combined MITE dataset is stored in SOURCEFORGE (https://sourceforge.net/projects/mite-mirna/files/). To comprehensively identify MITE-miRNAs in the analyzed species, 6,213 miRNAs were retrieved from PmiREN1.0 and 1,891 from miRBase after filtered with the latest criteria of plant miRNA annotation (Axtell and Meyers, 2018). The combined miRNA set was used to align against the identified MITEs by BLASTN (Camacho et al., 2009) and BLAT (Kent, 2002) with E-value < 1e^−10^. The miRNAs that contain over half sequences matched to MITEs were defined as MITE-miRNAs for further analysis.

### Identification of TID-miRNAs

A previously reported method (Allen et al., 2004) for identifying TID-miRNAs was employed with modifications on filtration. Briefly, genomic segments containing 500 base pairs at both flanking sides of miRNA precursors were extracted and used for sequence similarity search against the coding sequence (CDS) of all protein-coding genes using FASTA (v36) (Pearson and Lipman, 1988) and BLASTN (2.9.0+) (Camacho et al., 2009). After applying an E-value threshold of 1e^−10^, outputs by both aligning tools were combined and filtered with two stringent criteria. After removing hits with the matched sequences overlapping with miRNAs, only miRNAs contained two matched segments with opposite direction were considered as TID-miRNAs.

### Identification of LTR-miRNAs

The LTR retrotransposon sequences in *Z. mays* and *G. biloba* were obtained from the Plant Genome and Systems Biology Repeat Database (https://pgsb.helmholtz-muenchen.de/plant/recat/index.jsp) and GIGAdb (http://gigadb.org/dataset/100209), respectively. Full-length LTR retrotransposons in the other 18 species were first scanned by LTR_FINDER (v1.07) (Xu and Wang, 2007), and further filtered by LTR_retriever (v2.9.0) (Ou and Jiang, 2018). The resulting dataset for LTR retrotransposons is stored in SOURCEFORGE (https://sourceforge.net/projects/mite-mirna/files/). To identify candidate LTR-miRNAs, BLASTN (Camacho et al., 2009) was used to search sequence similarity between miRNA precursors and the identified LTR retrotransposons. Only those with E < 1e^−10^ and matched sequences longer than 50% of the precursors were considered LTR-miRNAs.

### Gene expression analysis in *O. sativa*

Thirteen RNA-Seq datasets for *O. sativa* ssp. *japonica* cv Nipponbare were downloaded from the Rice Genome Annotation Project (http://rice.plantbiology.msu.edu/expression.shtml) and used for quantifying gene expression. The data were processed using Tophat (Trapnell et al., 2009) and Cufflinks (Trapnell et al., 2010) to calculate the expression values in FPKM (mean fragments per kilobase per million mapped fragments) of annotated genes (MUS Release 7.0). Host genes of MITE-miRNAs and host genes of MITEs were identified using bedtools intersect (v2.29.2) (Quinlan, 2014) and their expression values were extracted for comparison.

### Quantification of miRNA expression by sRNA-Seq

The sRNA-Seq datasets employed to quantify miRNA expression were downloaded from NCBI SRA and listed in Supplementary Table 12. File uncompressing and adapter trimming were executed using the SRA toolkit and Trim Galore, respectively. Bowtie (1.1.2) (Langmead, 2010) was used to map the clean reads to the reference genomes allowing no mismatches. The expression values of mature miRNAs were normalized to RPM (reads per million) whereby the total numbers of uniquely mapped reads in the sRNA-Seq libraries were counted as the denominator.

### RT-qPCR analysis of miRNA abundance

Low-molecular-weight RNA was isolated from different tissues of *S. lycopersicum* cv Micro-Tom using the HiPure Plant miRNA Kit as recommended by the manufacturer (Magen). Reverse transcription of miRNAs into cDNAs was carried out using the miRcute Plus miRNA First-Stand cDNA Synthesis Kit (Tiangen). Quantitative real-time PCR was performed using the 7500 Fast Real-Time PCR engine (Applied Biosystems) and the miRcute Plus miRNA qPCR Kit with SYBR Green (Tiangen). Internal control for RT-qPCR was U6 (5’-TCGCTGAGCTTATTCTTTGTTCTG), and relative miRNA abundance was determined using the comparative threshold cycle method.

### Identification of MITE-miRNAs and nMITEs co-localized with pseudogenes

Annotation and genomic location of pseudogenes in *B. distachyon*, *G. max*, *M. truncatula*, *O. sativa*, and *S. bicolor* were obtained using the PseudoPipe workflow as previously reported (Xie et al., 2019). The obtained genomic locations were further extended by 2,000 base pairs to include the potential promoter regions. To obtain the genomic locations of MITEs, sequences of MITEs were aligned to reference genomes using minimap2 with default parameters (Li, 2018). The relative information is provided as Supplementary Table 13. Overlaps between the pseudogene loci and MITE-miRNAs and nMITEs were identified using bedtools intersect with parameter ‘-f 0.5’.

### Analysis of MITE features

MFE was calculated using RNAfold in the ViennaRNA package (version 2.2.10) (Denman, 1993). NMFE was determined through normalizing MFE against the length of MITEs. The distribution of NMFE in each species was plotted using a Python (version 3.6) script. For each MITE locus, the number of MITE family members, length, and GC content were calculated using in-house Perl scripts.

### XGBoost based machine learning model

To prepare the datasets for machine learning, four features of MITEs belonging to the *Mutator*, *PIF/Harbinger*, and *Tc1/Mariner* superfamilies in the 15 examined angiosperms were extracted as inputs. These MITEs were divided into mMITEs and nMITEs. A gradient boosting decision tree implemented in XGBoost (Extreme Gradient Boosting) (Chen and Guestrin, 2016) was used to construct a classifying model and evaluate importance of these features. The parameters were configured by default except the learning rate (eta=1) and maximum depth (max_depth=2). The objective function was set as “binary::logistic”, and boosting iteration number was set as 50. After the classifying model was created, the importance scores of the four features were calculated and ranked using the “xgb.importance” function in XGBoost.

### Identification and analysis of *Solyc12g008590* orthologous

Genome assemblies and annotation sequences of five species in *Solanum*, including *S. pimpinellifolium*, *S. pennellii*, *S. chilense*, *S. tuberosum*, and *S. melongena*, were downloaded from the Sol Genomics Network (https://solgenomics.net/). The Reciprocal Best Hits approach of BLASTP was used to identify orthologous genes of *Solyc12g008590* where the E value cutoff was set as 1e^−10^. The gene tree of *Solyc12g008590* homologs was obtained from Ensembl Plants (http://plants.ensembl.org/index.html).

### Target gene prediction

The psRNATarget program was used to predict miRNA targets using the V2 scoring schema (Dai et al., 2018). The expectation value was set to three instead of the default setting at five to keep only the high-confidence miRNA targets. All predicted target genes were listed in Supplementary Data 2.

### GO enrichment analysis

Annotations of GO terms were downloaded from Phytozome v11 (Goodstein et al., 2011). Association of the predicted target genes for MITE-miRNAs, LTR-miRNAs, and TID-miRNAs with the GO terms was analyzed using customized scripts. Fisher’s exact test with the Benjamini-Hochberg correction was used to find enriched GO terms with the adjusted *P* value set at 0.05.

### Polymorphisms of miRN2285 and its target gene

Polymorphisms of Osa-miRN2285 and its target gene, *LOC_Os06g39750*, and phenotypic records on cold tolerance (BPCT) of different strains were retrieved from MBKBASE (Peng et al., 2019) (http://mbkbase.org/). Statistical analysis was done through the independent samples t-test.

### Association of summer temperature with *O. sativa* germplasm distribution

Geographical information of *O*. *sativa* germplasms in China, including longitude and latitude, was retrieved from MBKBASE and shown in Supplementary Table 14. Temperature data in China from 2009 to 2018 were downloaded from the National Earth System Science Data Center, National Science & Technology Infrastructure of China (http://www.geodata.cn). The mean of summer temperature in June, July and August were calculated by a customized MATLIB R2019b (9.7.0.1190202) script.

### Other statistical analyses

If not stated specifically, the comparison of two distributions of values was tested with an independent samples t-test. *P* values were shown as exact values or otherwise referenced with a symbol according to the following scales: *, *P* < 0.05; **, *P* < 0.01.

## Code availability

All codes used in this study are freely accessible via the GitHub repository (https://github.com/little-raccoon/MITE-miRNA/).

## Acknowledgments

We are indebted to Hailong Chen for assistance with obtaining the temperature data. This work was supported by grants from the National Key Research and Development Program of China (2018YFE0204700) and the National Natural Science Foundation of China (31621001). Part of the bioinformatics analysis was performed on the High-Performance Computing Platform of the Center for Life Science at Peking University.

## Ethics declarations

The authors declare no declare no conflict of interest.

---

**Supplementary Fig. 1.**
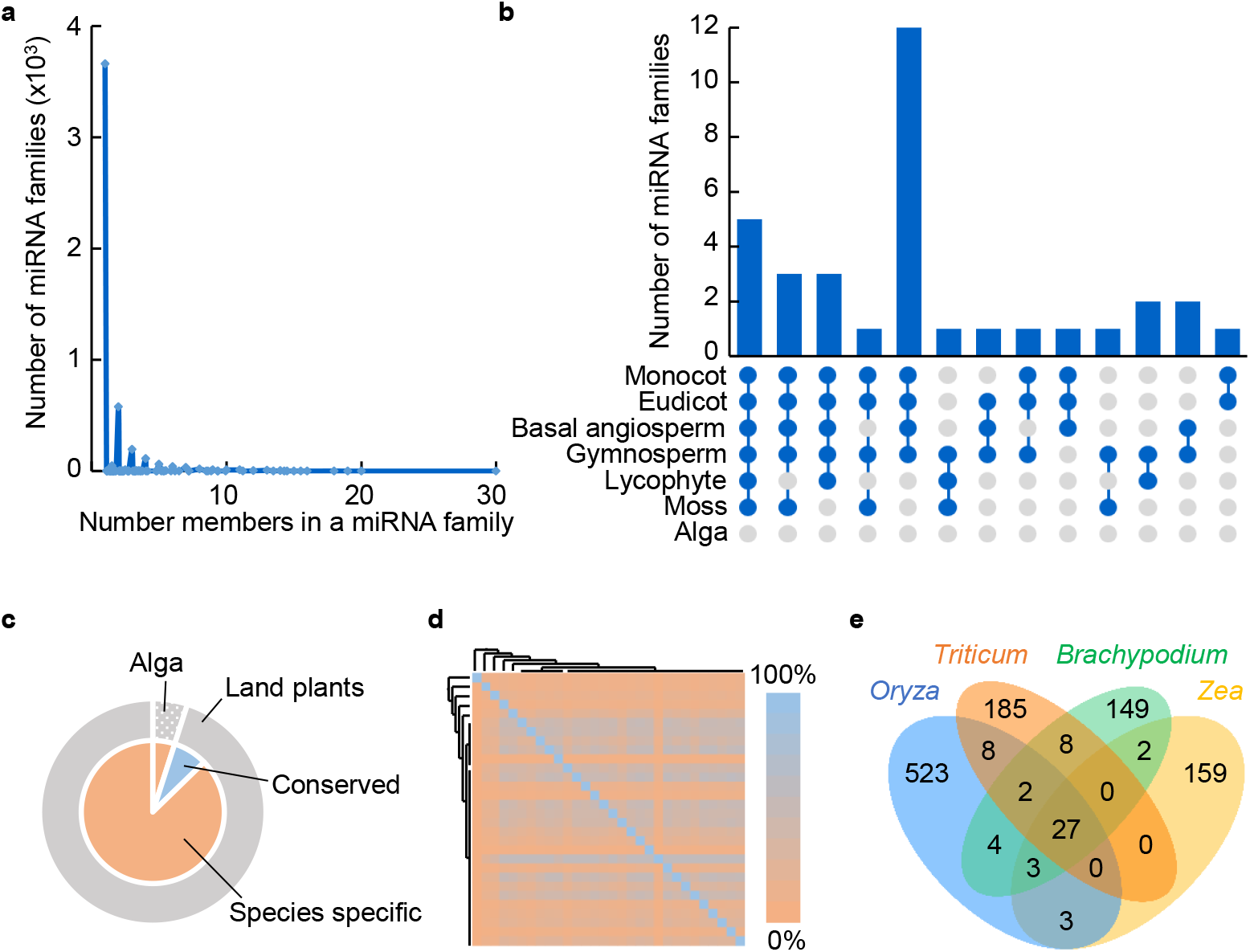
Analysis of miRNA family conservation in selected plants. **a**, Plotting the number of miRNA families and the number members in the miRNA families. Annotated miRNAs in 88 different plant species were grouped into families based on similarity of the mature miRNA. **b**, Number of miRNA families conserved at different taxonomic levels. The 88 plant species were divided into seven taxonomic groups. Blue dots indicate a given miRNA family is detected in over half species in a taxonomic group. Histograms show the total number of miRNA families corresponding to different connecting patterns of the dots. **c**, Proportions of conserved and species-specific miRNA families in examined green algae and land plants. **d**, Pairwise comparison for the conservation of miRNA families among 30 taxonomic families. Color of the heatmap represents the proportion of miRNA families in a given taxonomic family that are detected in another family. **e**, Venn diagram showing the number of miRNA families detected in the four examined genera of the Poaceae family.

**Supplementary Fig. 2.**
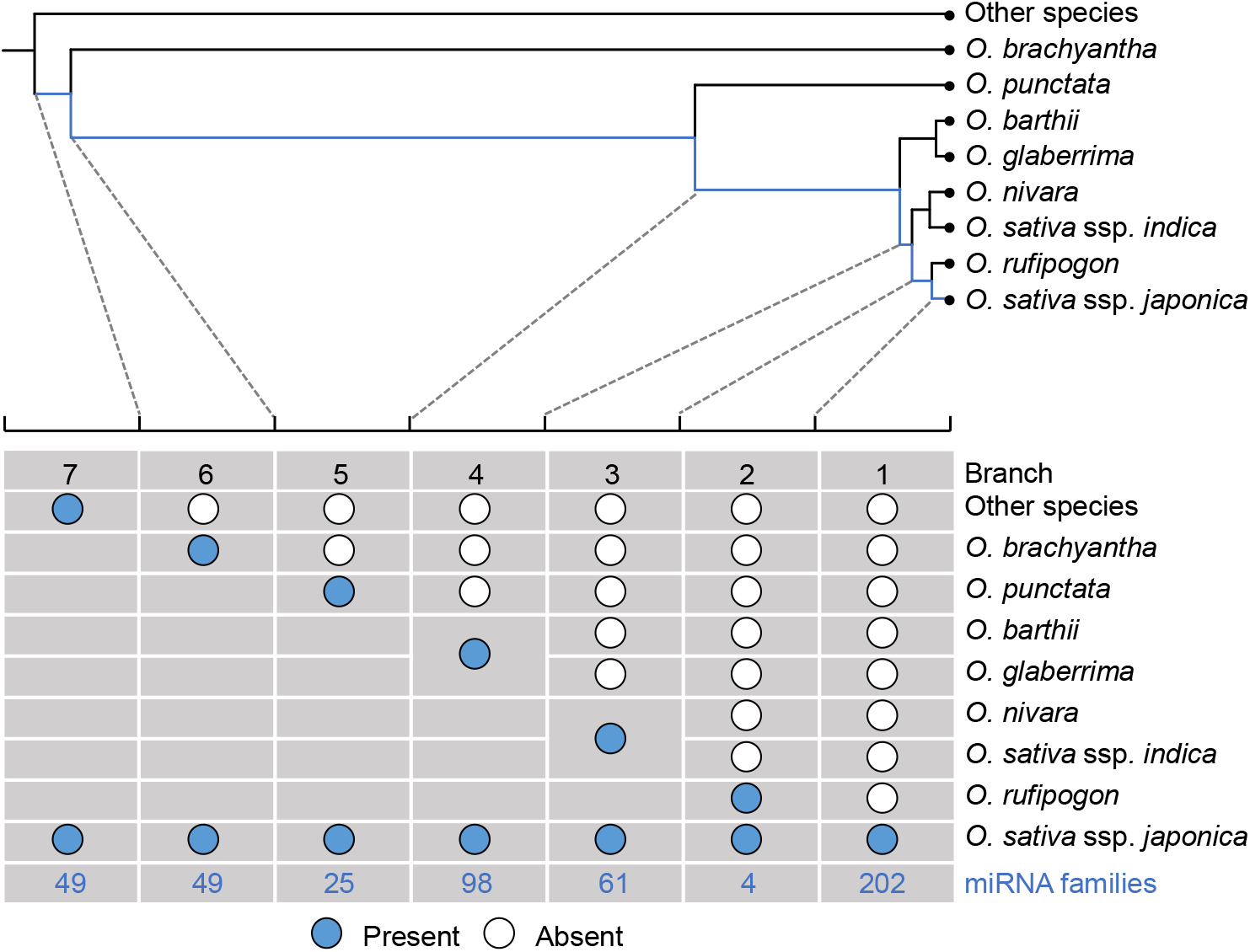
Rationale for identifying *de novo* miRNA families in *Oryza*. Origination events for novel miRNA families in *O. sativa* ssp. *japonica* were identified by comparing the miRNA portfolios in eight species or subspecies in the genus of *Oryza*. A parsimony rule was applied to assign the ancestral or derivative states of a miRNA family in *O. sativa* ssp. *japonica*. According to the phylogenetic tree of *Oryza*, stepwise comparisons were made to identify novel miRNA families in *O. sativa* ssp. *japonica* that were absence in a diverged species but presence in a species sharing an ancestor with *O. sativa* ssp. *japonica* after the divergence. Numbers of miRNA families identified at each branch are indicated at bottom. The phylogenetic tree and estimated divergence time in each internal node were retrieved from the TimeTree database.

**Supplementary Fig. 3.**
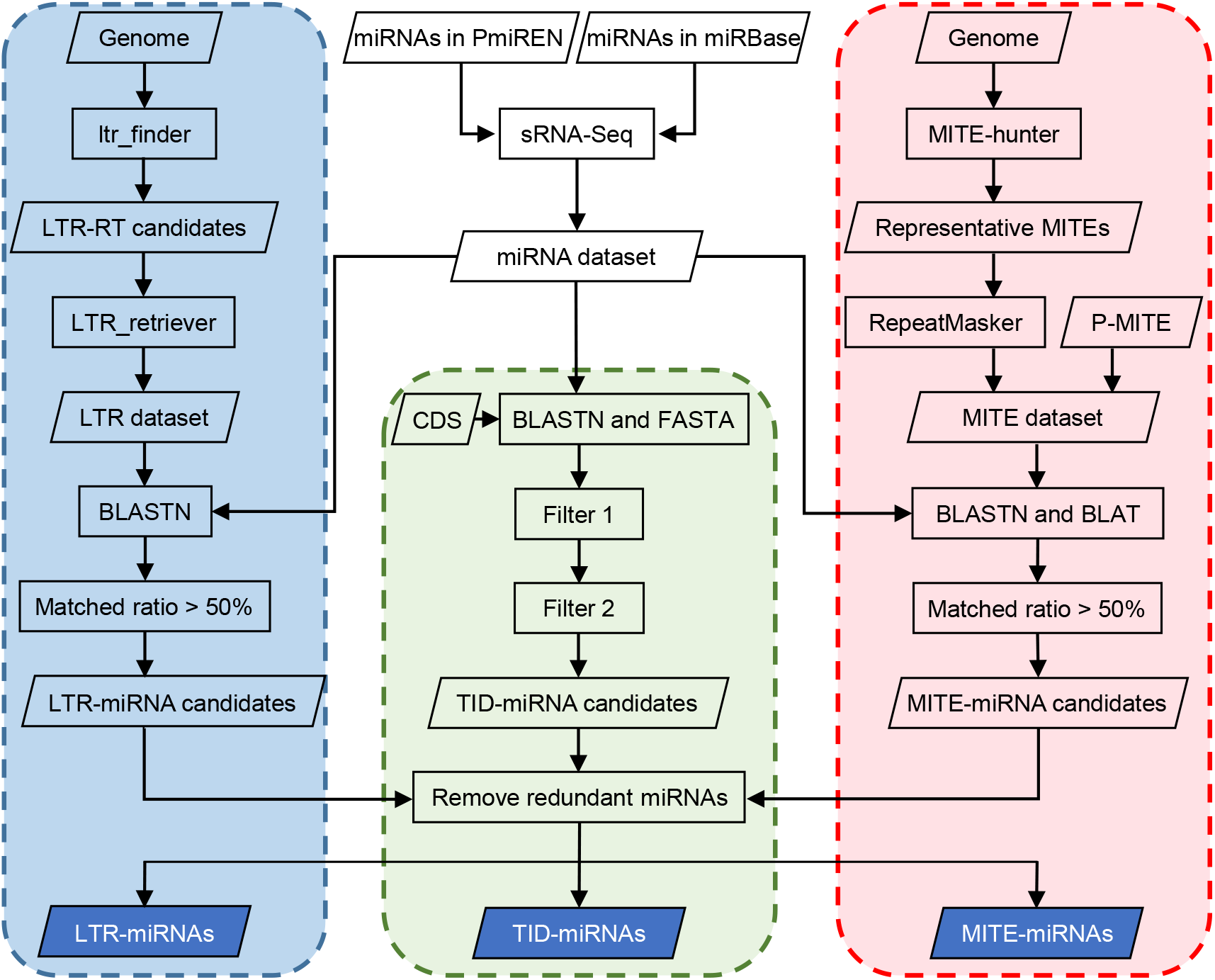
Workflow for detecting the three types of de novo miRNAs. Twenty phylogenetically representative species with high-quality reference genomes (genome) and available small RNA sequencing data (sRNA-Seq) were used to identify the three types of novel miRNAs. Computational pipelines for detecting LTR-miRNAs, TID-miRNAs, and MITE-miRNAs are shown from left to right. In identifying TID-miRNAs, two filters were applied. First, when the matched sequences overlapped with a known miRNA, the hits were removed. Second, only candidates containing two matched segments with opposite directions were further processed as candidate TID-miRNAs. In the rare cases when a miRNA was detected by more than one pipeline, the finial assignment was determined by the longest match of the miRNA locus against sequences of a MITE, an LTR, or a protein-coding gene to remove redundancy.

**Supplementary Fig. 4.**
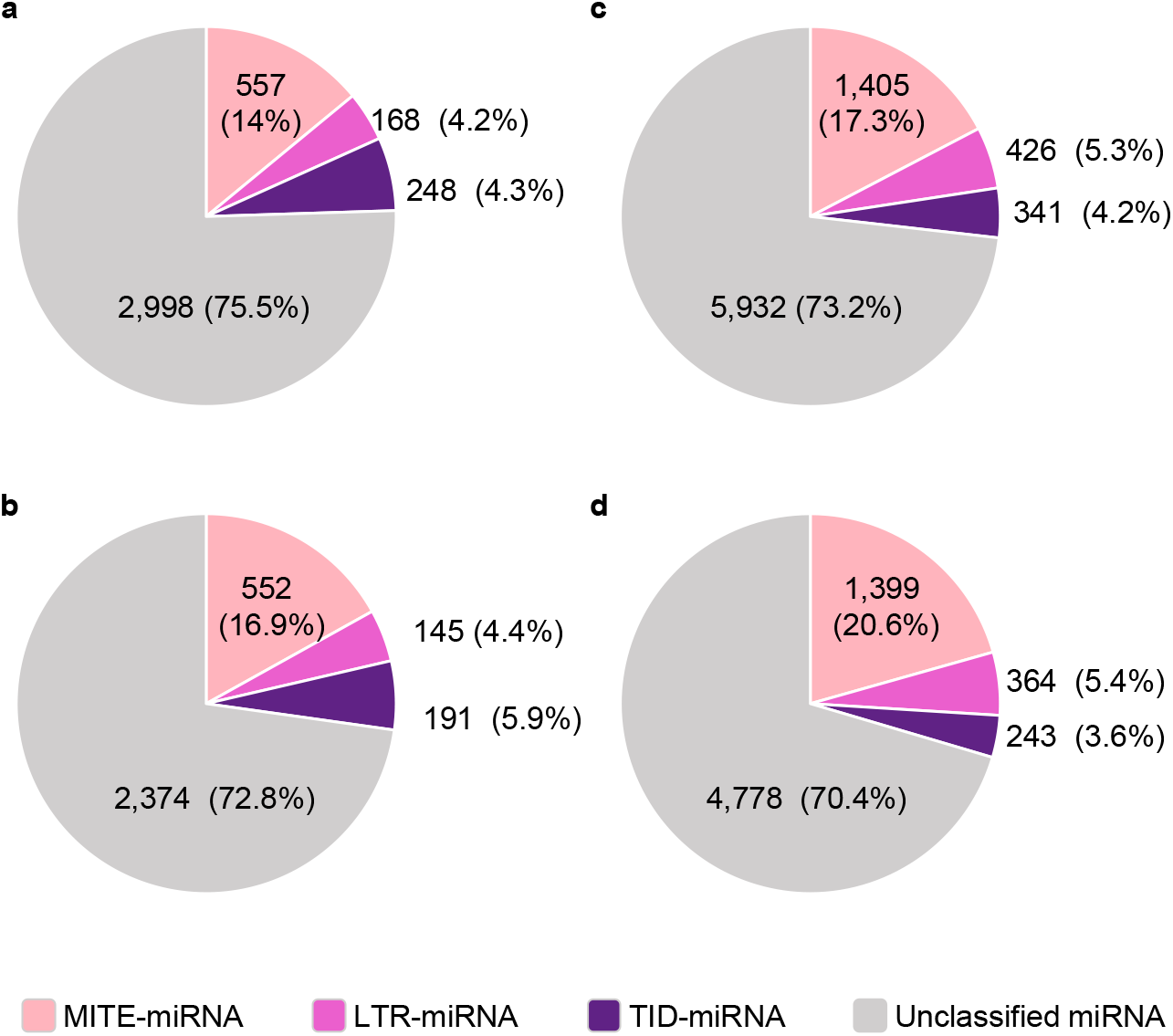
MITE-miRNAs are the dominant *de novo* miRNAs in plants. **a-b**, Pie graphs showing the proportions of MITE-miRNA families, LTR-miRNA families, TID-miRNA families, and unclassified miRNA families in the 20 examined plants (**a**) and the 15 examined angiosperms (**b**). **c-d**, Proportions of MITE-miRNAs, LTR-miRNAs, TID-miRNAs and unclassified miRNAs in the 20 plants (**c**) and the 15 angiosperms (**d**).

**Supplementary Fig. 5.**
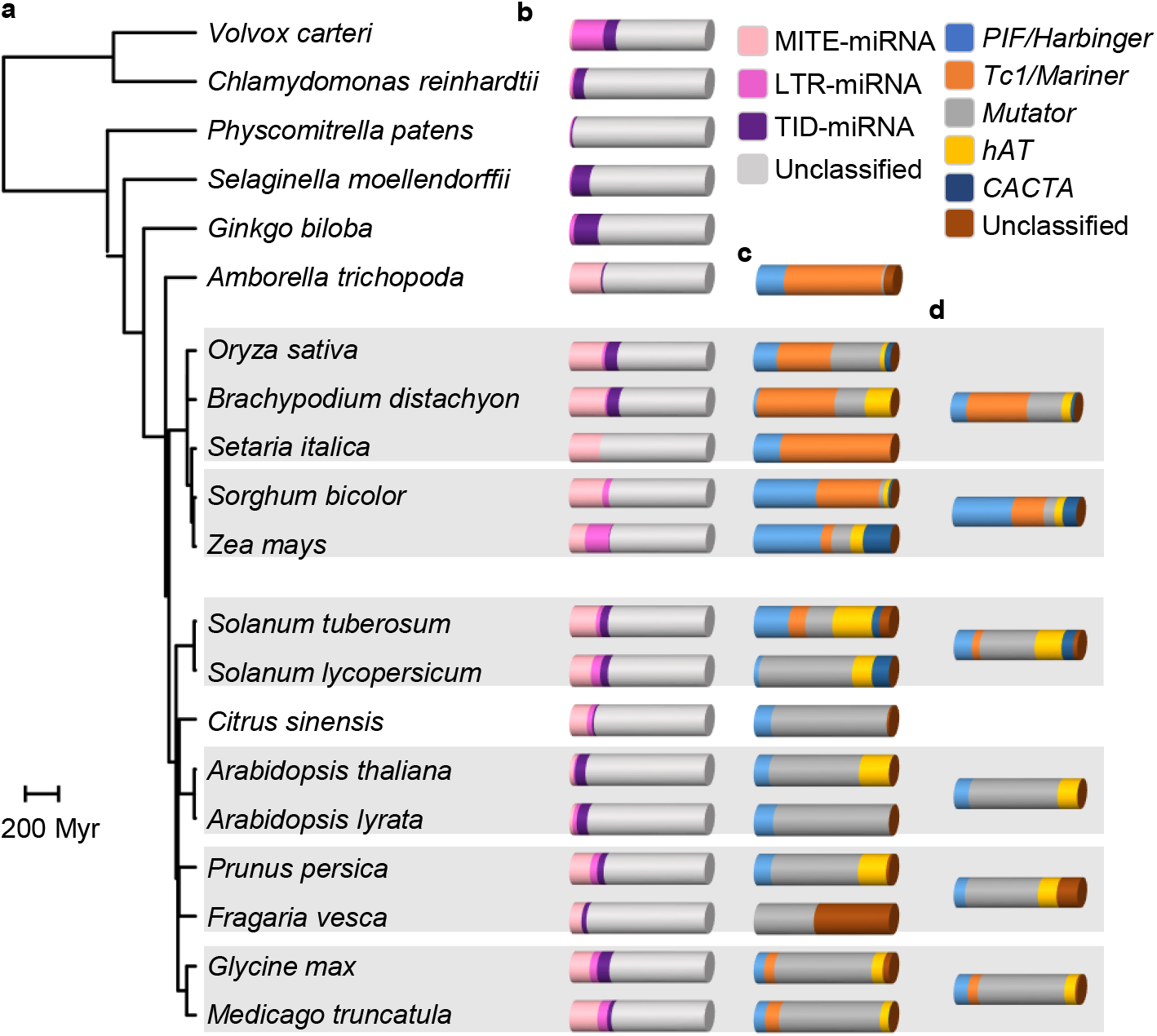
Bursts of MITE-miRNAs in angiosperm. **a**, Phylogenetic tree of the 20 examined plant species retrieved from TimeTree. **b**, Bar graphs showing proportions of MITE-miRNAs, LTR-miRNAs, TID-miRNAs, and unclassified miRNAs in each species. The proportions were calculated by dividing the numbers of identified MITE-miRNAs, LTR-miRNAs, TID-miRNAs, and the remaining miRNAs against the total numbers of miRNA in a species. **c**, Proportion of MITE-miRNAs derived from the five MITE superfamilies in individual species. **d**, Average proportion of MITE-miRNAs derived from the five MITE superfamilies in the six taxonomic groups as shaded in (**a**).

**Supplementary Fig. 6.**
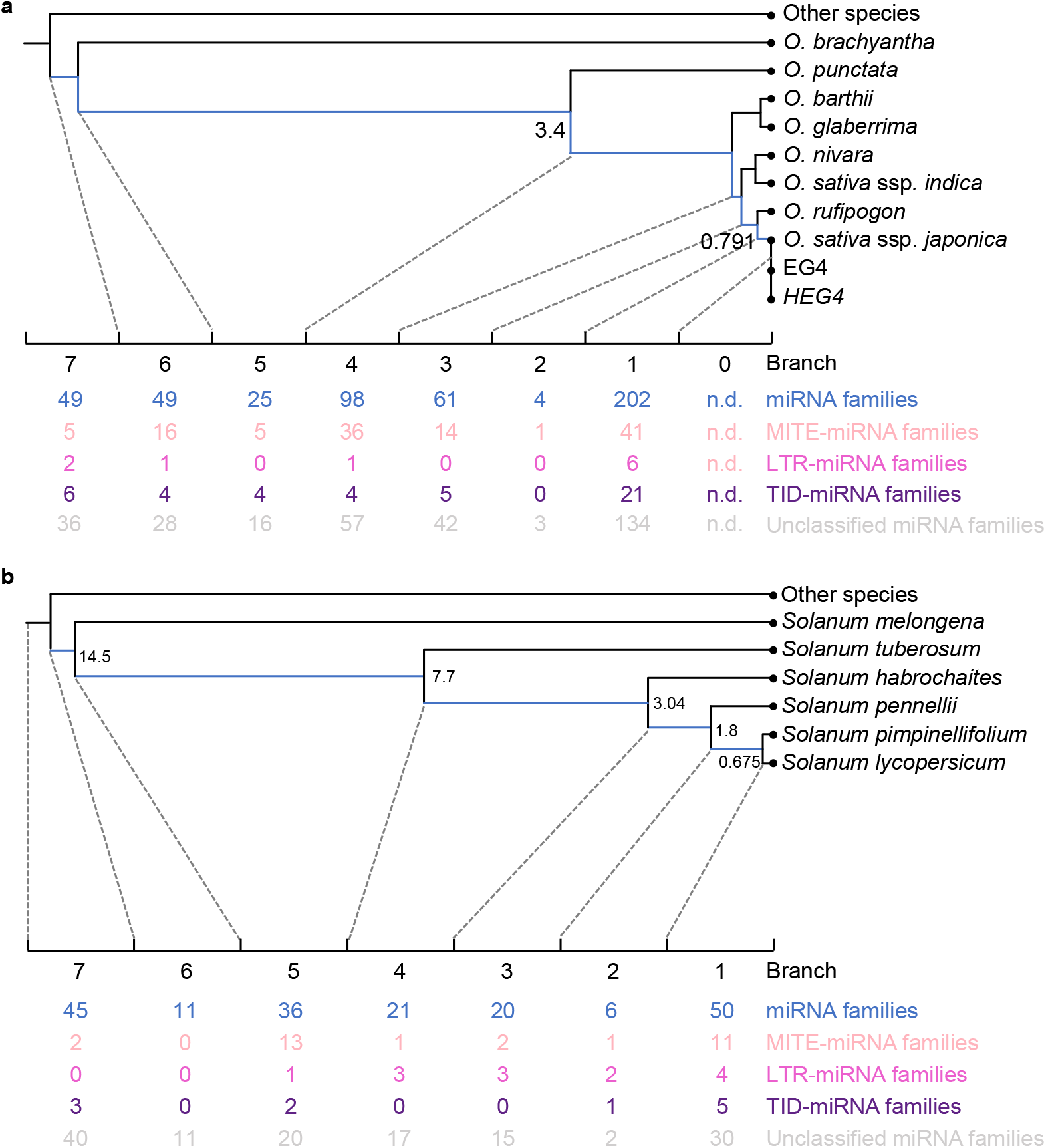
Evolutionary distribution of the origination events for the three types of *de novo* miRNA families in the *Oryza* and *Solanum* genera. **a-b**, Evolutionary distribution of origination events for *de novo* miRNA families in *Oryza* towards *O. sativa* ssp. *japonica* (**a**) and in *Solanum* towards *S. lycopersicum* (**b**). The phylogenetic trees and estimated divergence time were retrieved from TimeTree. The numbers indicate novel MITE-miRNA families, LTR-miRNA families, TID-miRNA families, and unclassified miRNA families that were identified in each branch. n.d., not determined.

**Supplementary Fig. 7.**
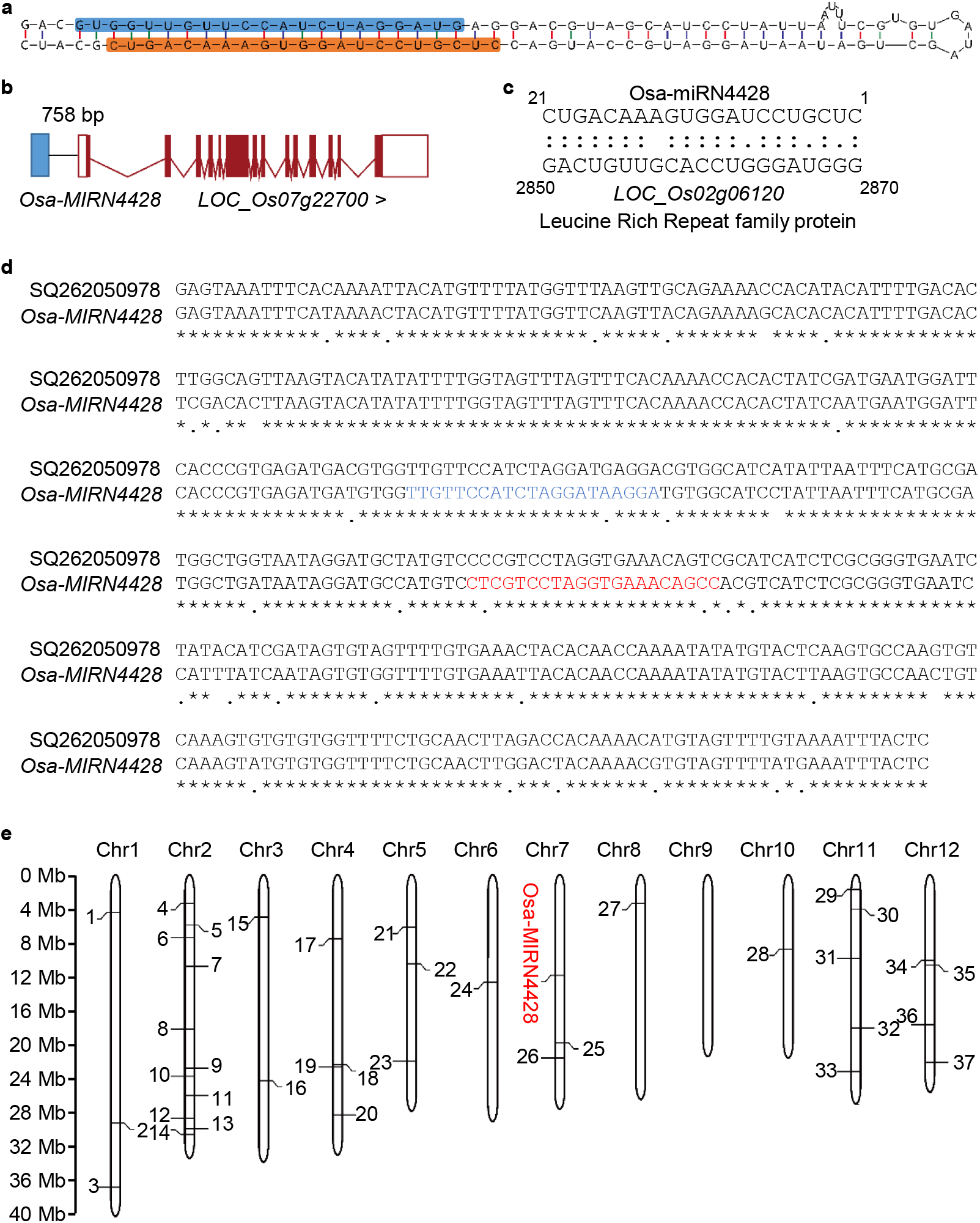
Characterization of the MITE-miRNA Osa-miRN4428. **a**, Predicted secondary structure of the *Osa-MIRN4428* precursor. The mature and star sequences of Osa-miRN4428 were highlighted in orange and blue, respectively. **b**, *Osa-MIRN4428* is located near the gene *LOC_Os07g22700*, only 758 bp upstream of its transcriptional start site. **c**, Complementarity pattern between Osa-miRN4428 and its predicted target site in *LOC_Os02g06120*, which encodes a Leucine Rich Repeat family protein. Perfect and wobble base pairs are indicated by “:” and “.”, respectively. **d**, Sequence alignment between *Osa-MIRN4428* and a homologous *Mutator* MITE (SQ262050978). Mature and star sequences of Osa-miRN4428 are shaded in red and blue, respectively. **e**, Chromosomal locations of *Osa-MIRN4428* and 37 homologous MITEs in the *O*. *sativa* genome.

**Supplementary Fig. 8.**
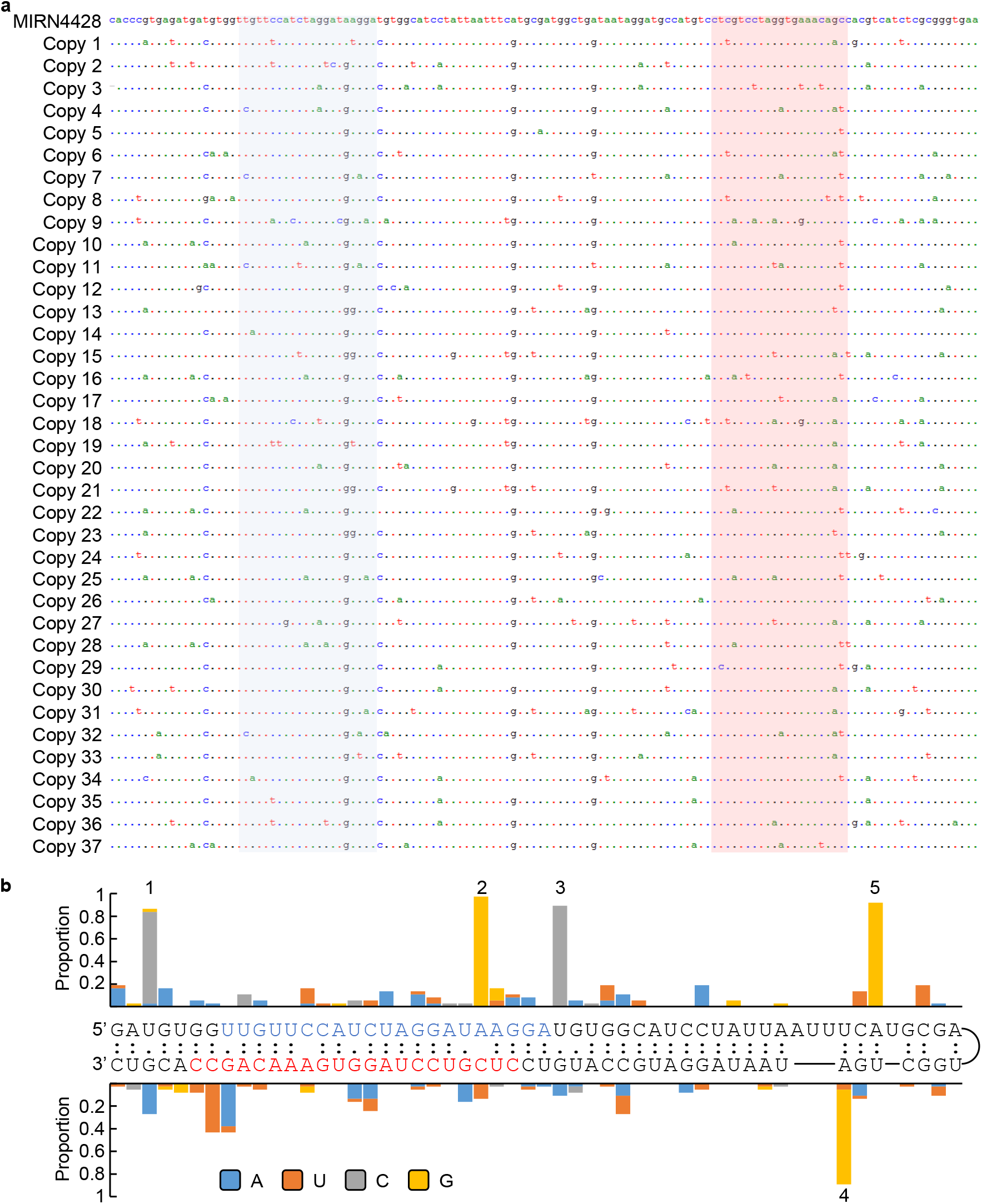
Sequence comparison among *Osa-MIRN4428* and 37 related MITEs. **a**, Multiple sequence alignment of *Osa-MIRN4428* and 37 copies belonging to the same SQ262050978 MITE family. Mature and star sequences of Osa-miRN4428 are shaded in red and blue, respectively. **b**, Distribution of SNPs along the *Osa-MIRN4428* precursor. Histogram showing the proportion of SNPs on the *Osa-MIR4428* precursor. Base composition is shown in different color. Secondary structure of the *Osa-MIRN4428* precursor was predicted by RNAfold. Perfect and wobble base pairs are indicated by “:” and “.”, respectively. The mature and star sequences of Osa-miRN4428 are highlighted in red and blue, respectively. Five major SNPs are marked with numbers from left to right.

**Supplementary Fig. 9.**
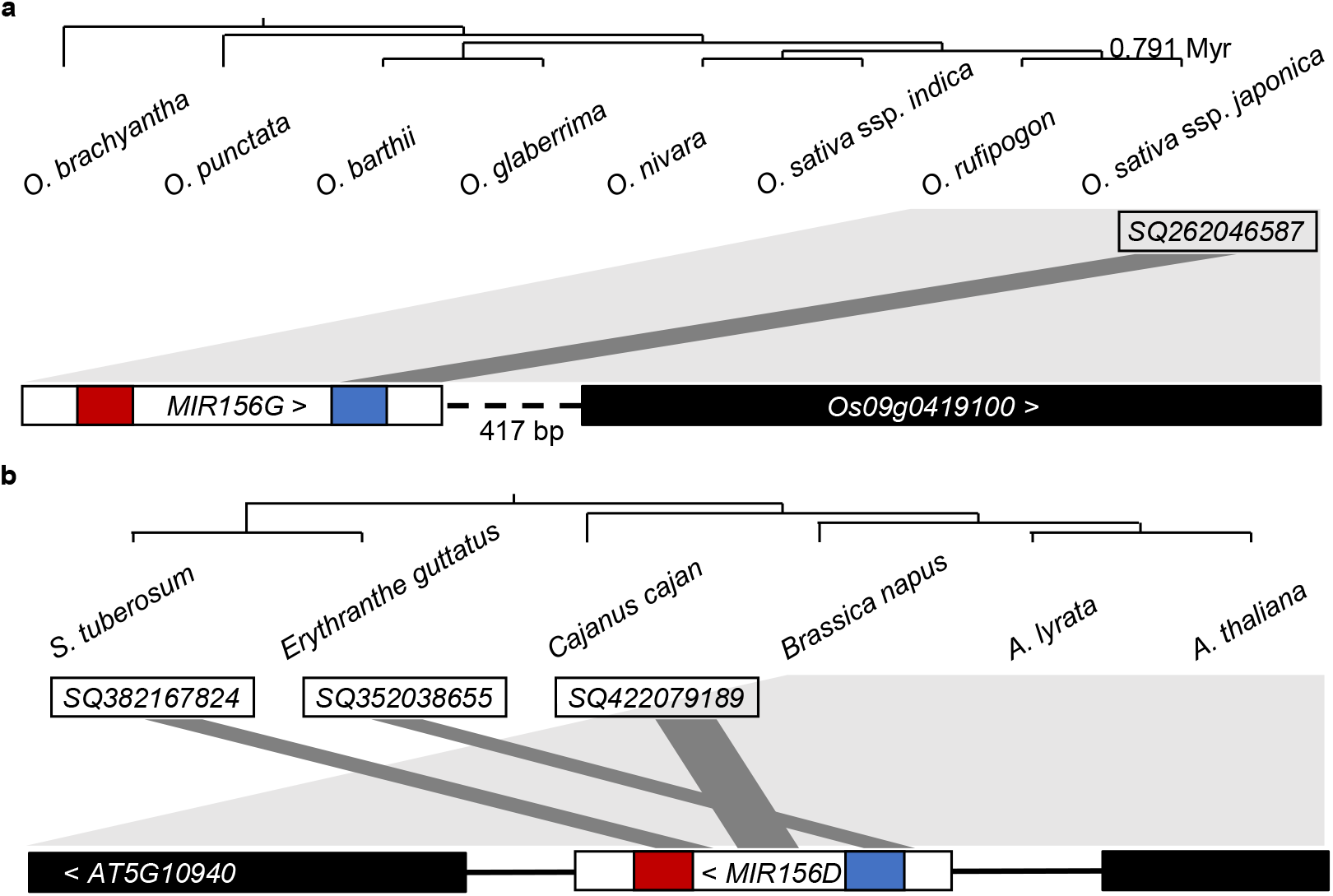
Synteny analyses of two putative MITE-miRNAs belonging to the conserved miR156 family. **a-b,** Synteny analyses of the *Ath-MIR156D*-containing fragment (**a**) and the *Osa-MIR156G*-containing fragment (**b**). The phylogenetic trees consisting of relative species were retrieved from TimeTree.org. The syntenic blocks are shown at the bottom with exons depicted as boxes, introns as horizontal lines, and promoters as dashed lines. Arrows mark transcription directions of the genes. Regions corresponding to the mature and star miRNAs are represented by red and blue squares, respectively. Species containing the synteny block are shaded with light grey. Dark grey binds indicate sequences shared between the miRNAs and the MITEs in the given species, which are depicted as open boxes.

**Supplementary Fig. 10.**
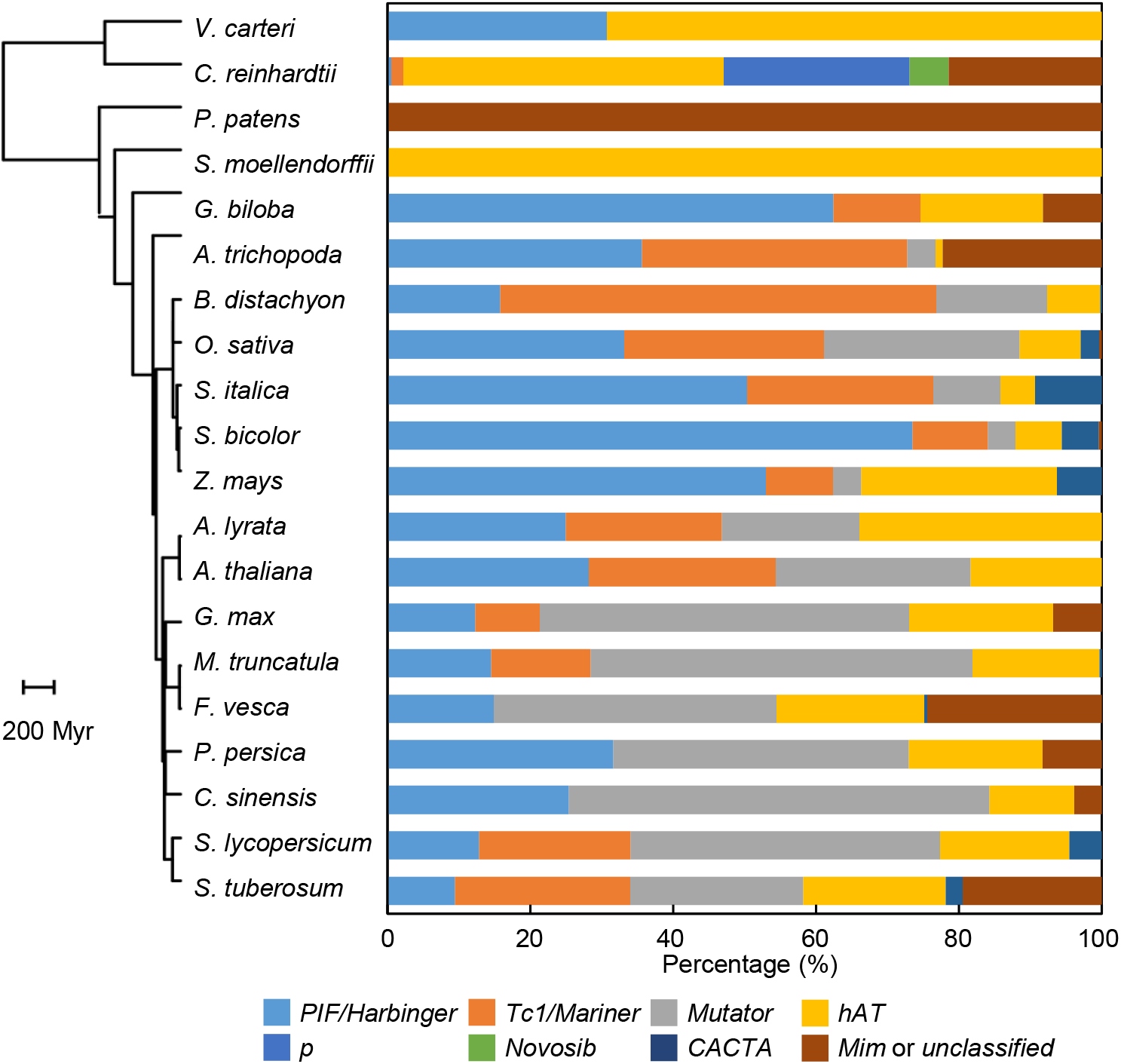
MITE constitution in 20 examined plants. The 3,070,716 identified MITEs in 19 examined species were divided into seven superfamilies based on homology of the TIR and TSD. MITEs in *P*. *patens* were not classified into superfamilies due to ambiguous TSD and/or TIR features (Chen et al., 2014). Proportions of MITEs belonging to the seven superfamilies and not assigned to any superfamily (unclassified) are shown next to each species, which are grouped into a phylogenetic tree retrieved from the TimeTree database.

**Supplementary Fig. 11.**
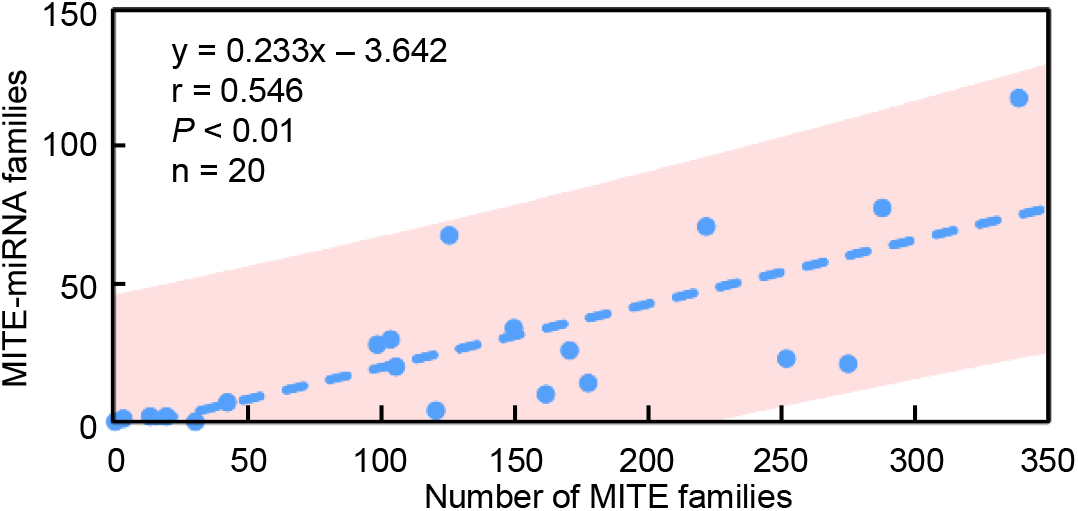
Linear regression analysis between the number of MITE families and the number of MITE-miRNA families in the 20 examined plants. Pearson’s correlation was used to determine the linear relationship between the number of MITE families and the number of MITE-miRNA families in the 20 examined plants. Dotted blue line represents the deduced linear regression (r = 0.546, *P* < 0.01) with the 95% confidence intervals shaded in pink.

**Supplementary Fig. 12.**
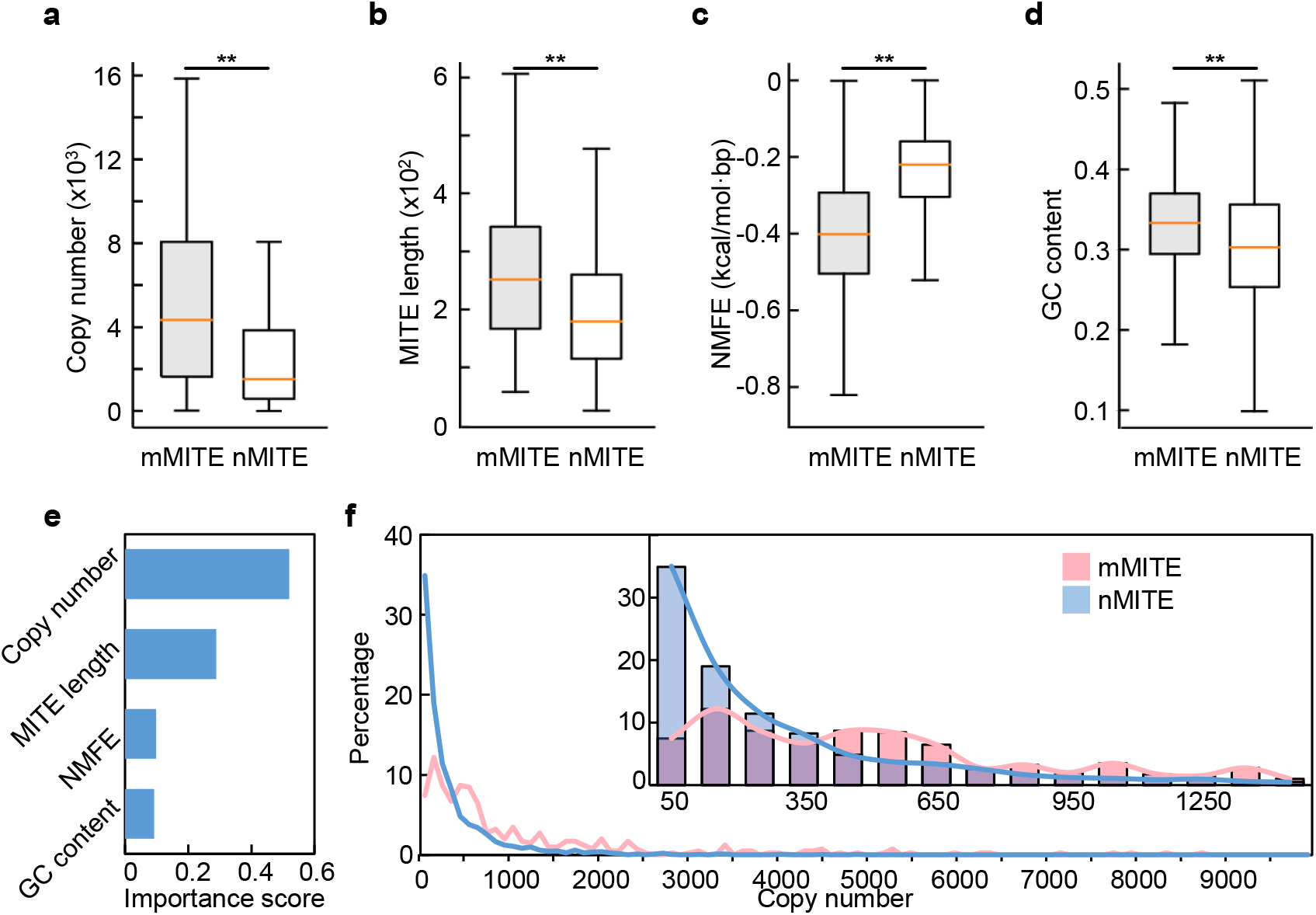
MITE features influencing conversion to miRNAs. **a-d**, Comparison of mMITEs and nMITEs regarding copy number of the MITE family members (**a**), length (**b**), NMFE (**c**), and GC content (**d**) of the MITEs. **, *P* < 0.01 by independent samples t-test. **e**, Assessment for relative importance of the four examined features by a machine learning approach based on a gradient boosting decision tree algorithm. **f**, Comparison of the copy number of mMITE and nMITE families in the 15 examined angiosperms. Line chart shows the proportions of MITE families with a given copy number. The line chart and histogram in the inlet show the length distribution of MITE families with copy number below 1,500. The x-axis represents the bin median where the bin size is 100.

**Supplementary Fig. 13.**
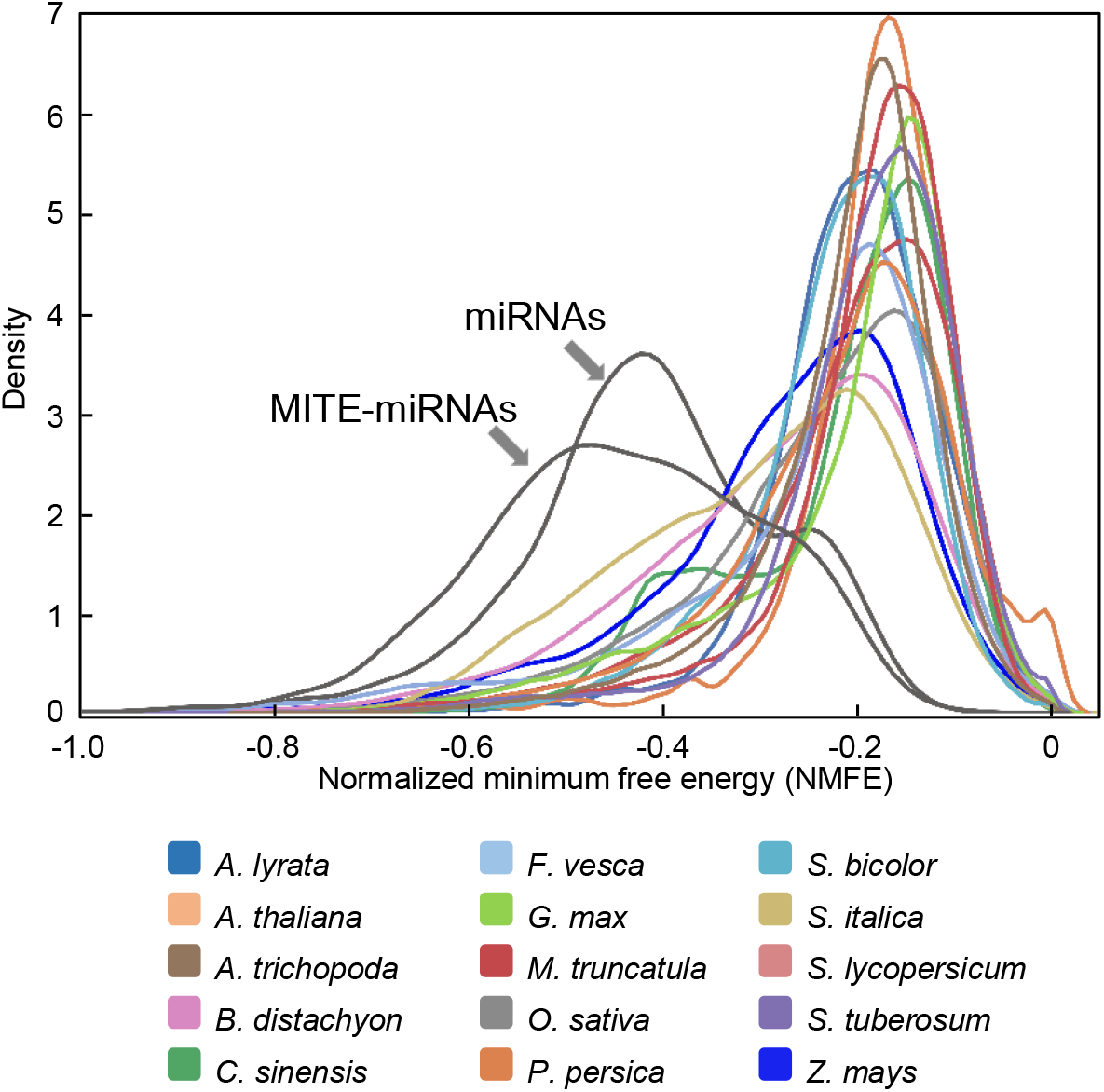
Comparison of the NMFE between miRNAs and MITEs. Minimal free energy for all miRNAs, MITE-miRNAs, and MITEs in the 15 examined angiosperms was calculated using RNAfold in the ViennaRNA package (version 2.2.10). NMFE was calculated by dividing minimal free energy by the sequence length. Kernel density estimation was used to display the distribution of NMFE. NMFE distribution of MITEs in the 15 species was shown in different colors.

**Supplementary Fig. 14.**
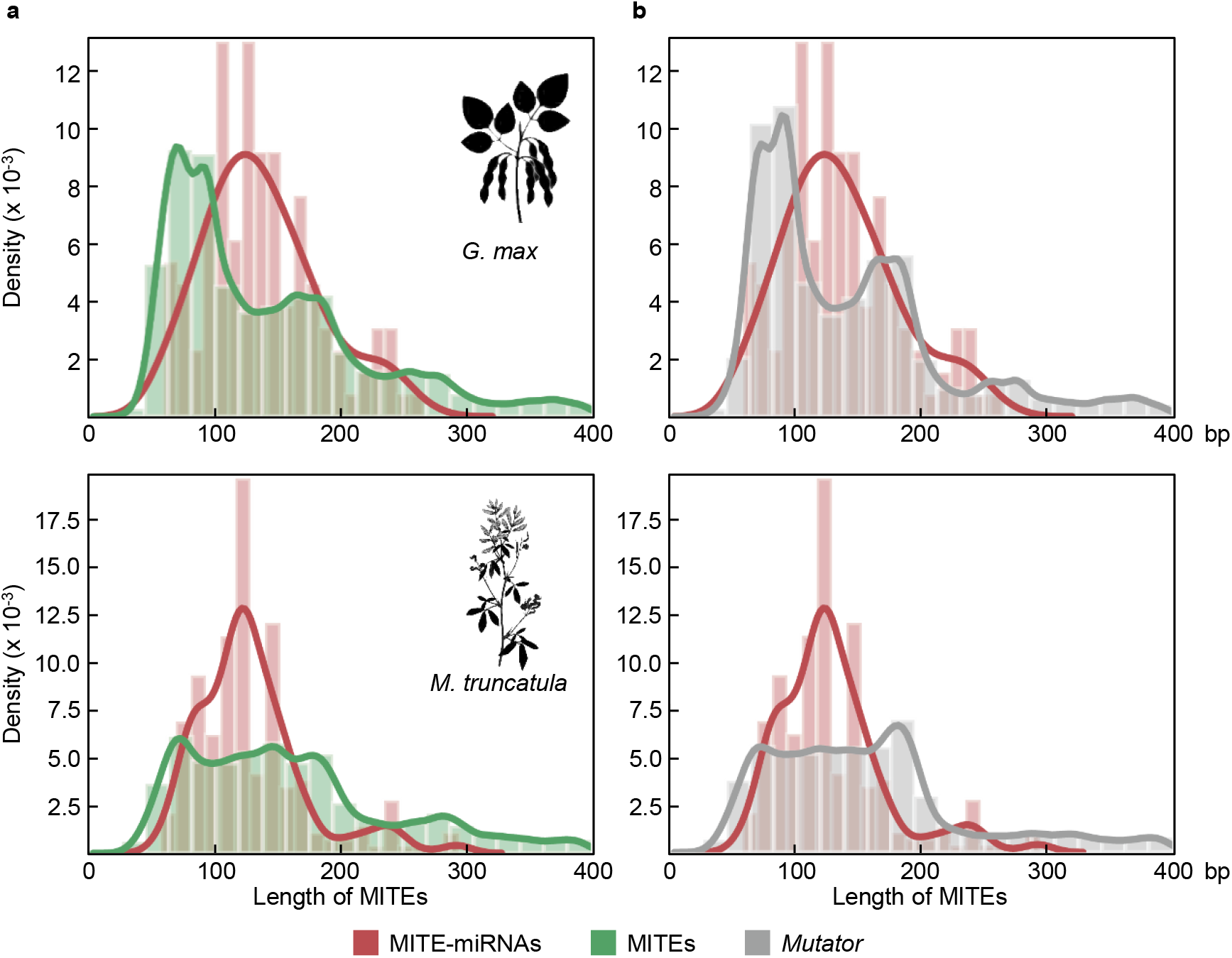
Comparison of length distribution of MITE-miRNAs and MITEs in *G. max* and *M. trucatula.* **a**, Length of all MITE-miRNAs and MITEs in *G. max* (top) and *M. truncatula* (bottom) was calculated. Length distribution was determined using kernel density estimation with histograms. **b**, Comparison of the length distribution of MITE-miRNAs and MITEs of the *Mutator* superfamily in *G. max* and *M. truncatula*.

**Supplementary Fig. 15.**
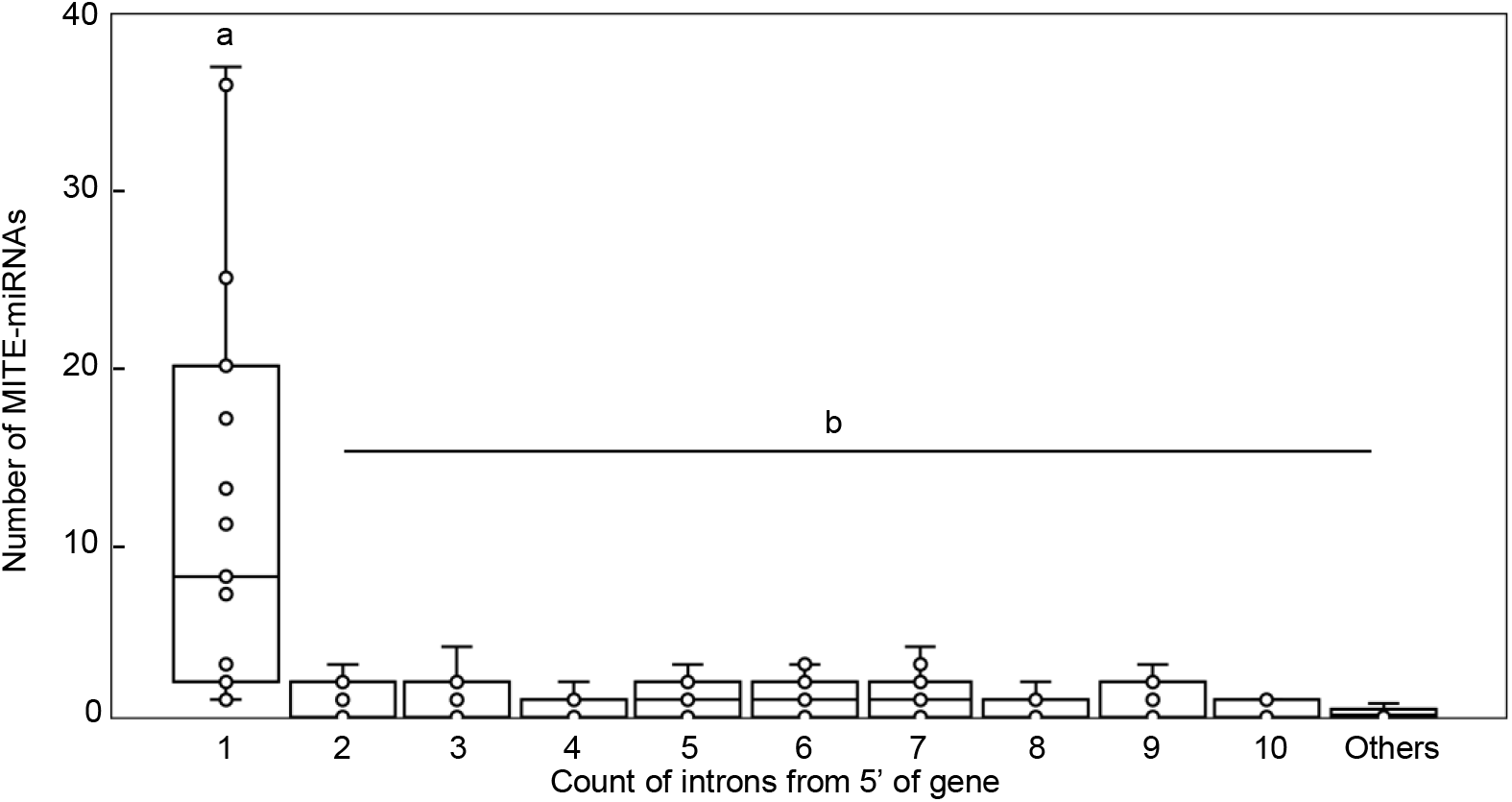
Preferential localization of MITE-miRNAs in the first intron. Number of MITE-miRNAs in different introns was calculated and plotted against the order of the introns counting from the 5’ end of the gene. The box represents values corresponding to the 25^th^ and 75^th^ percentile. Horizontal line inside the box represents the median. Different letters denote groups with significant differences (*P* < 0.01 by independent samples t-test).

**Supplementary Fig. 16.**
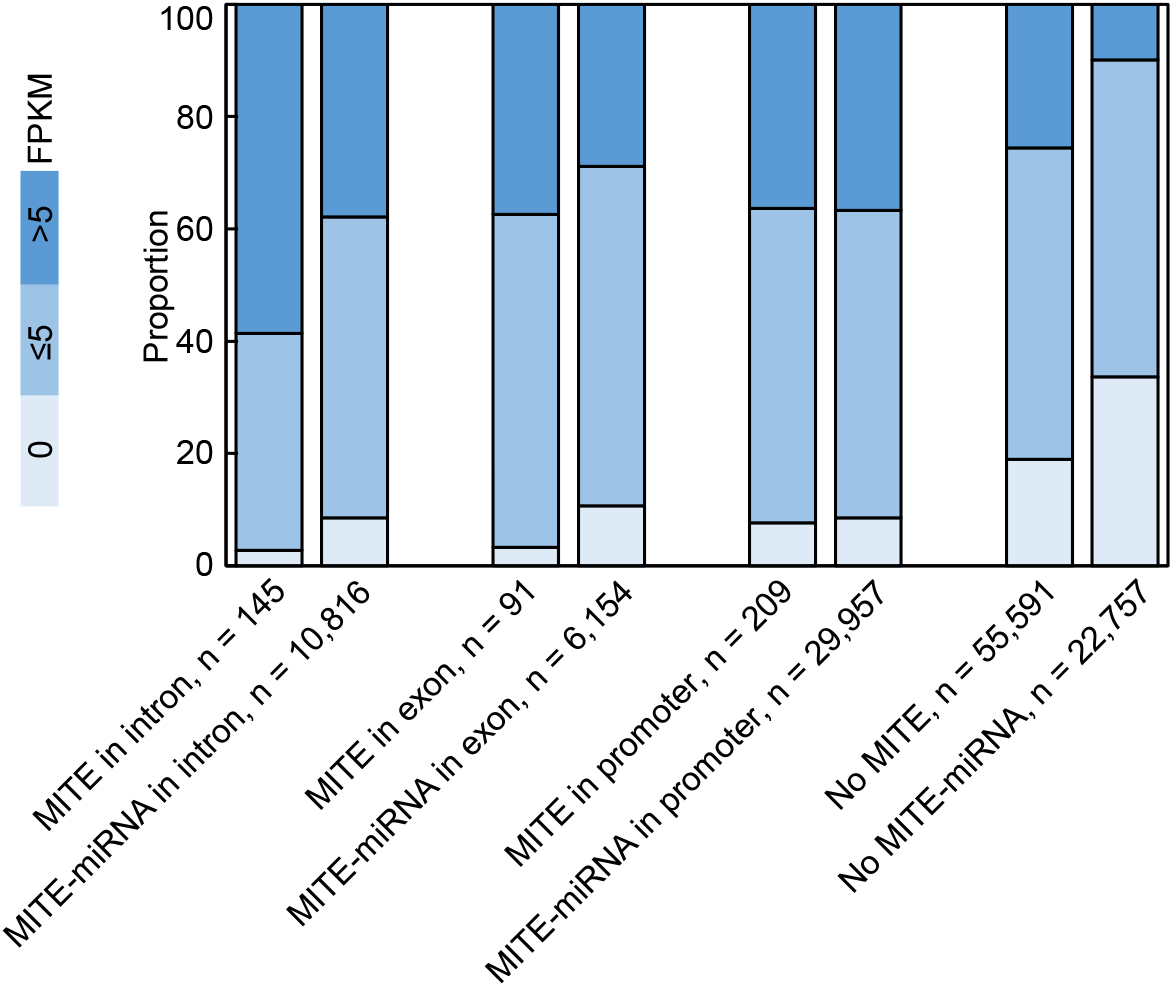
Comparison of the expression level of MITE-miRNA host genes and MITE host genes in *O. sativa*. Annotated genes were divided into different groups based on whether they contain a MITE or a MITE-miRNA and where the MITE or MITE-miRNA locates. Gene expression level was determined by the fragments per kilobase per million mapped fragments method. Values are the average of 13 RNA-Seq datasets with the accession numbers SRX100741, SRX100743, SRX100745, SRX100746, SRX100747, SRX100749, SRX100753, SRX100754, SRX100755, SRX100756, SRX100757, SRR042529, and SRX016110.

**Supplementary Fig. 17.**
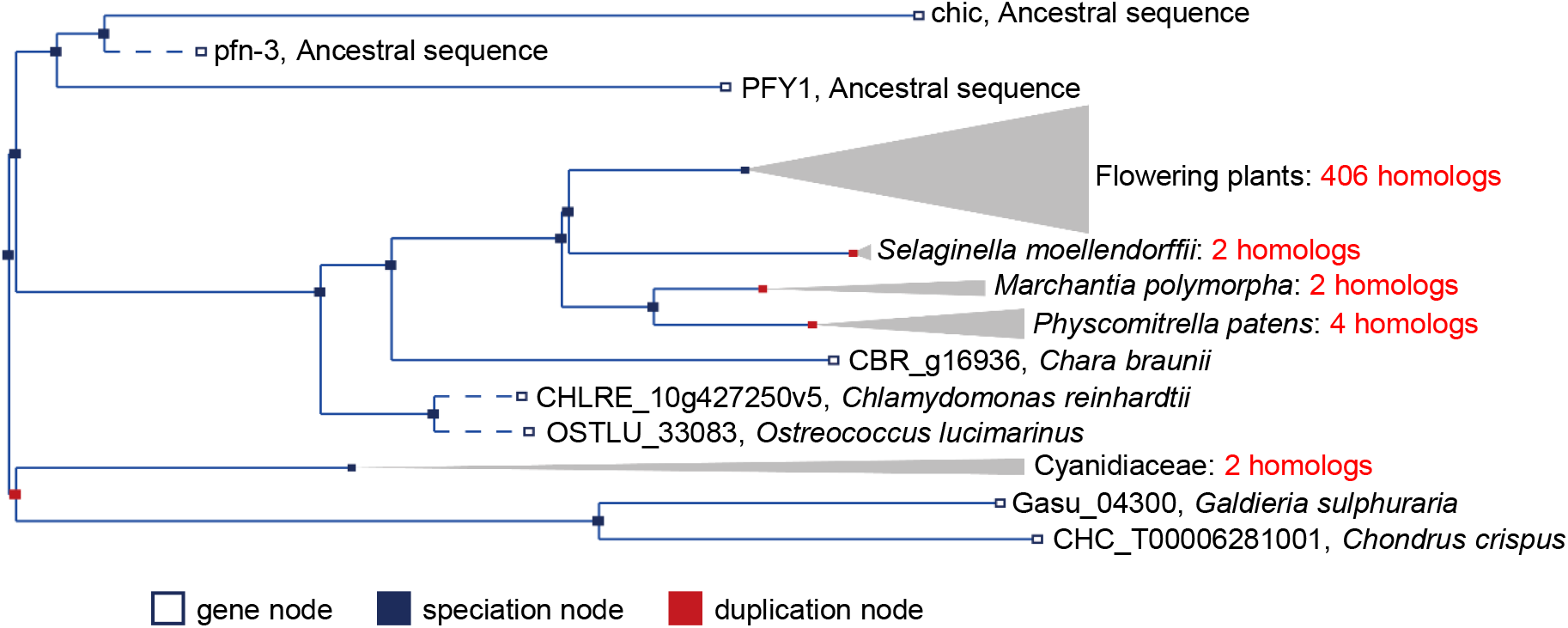
Gene tree representing the *Solyc12g008590* orthologues. The tree was generated using the Gene Orthology/Paralogy prediction method pipeline in EnsemblPlants (https://plants.ensembl.org).

**Supplementary Fig. 18.**
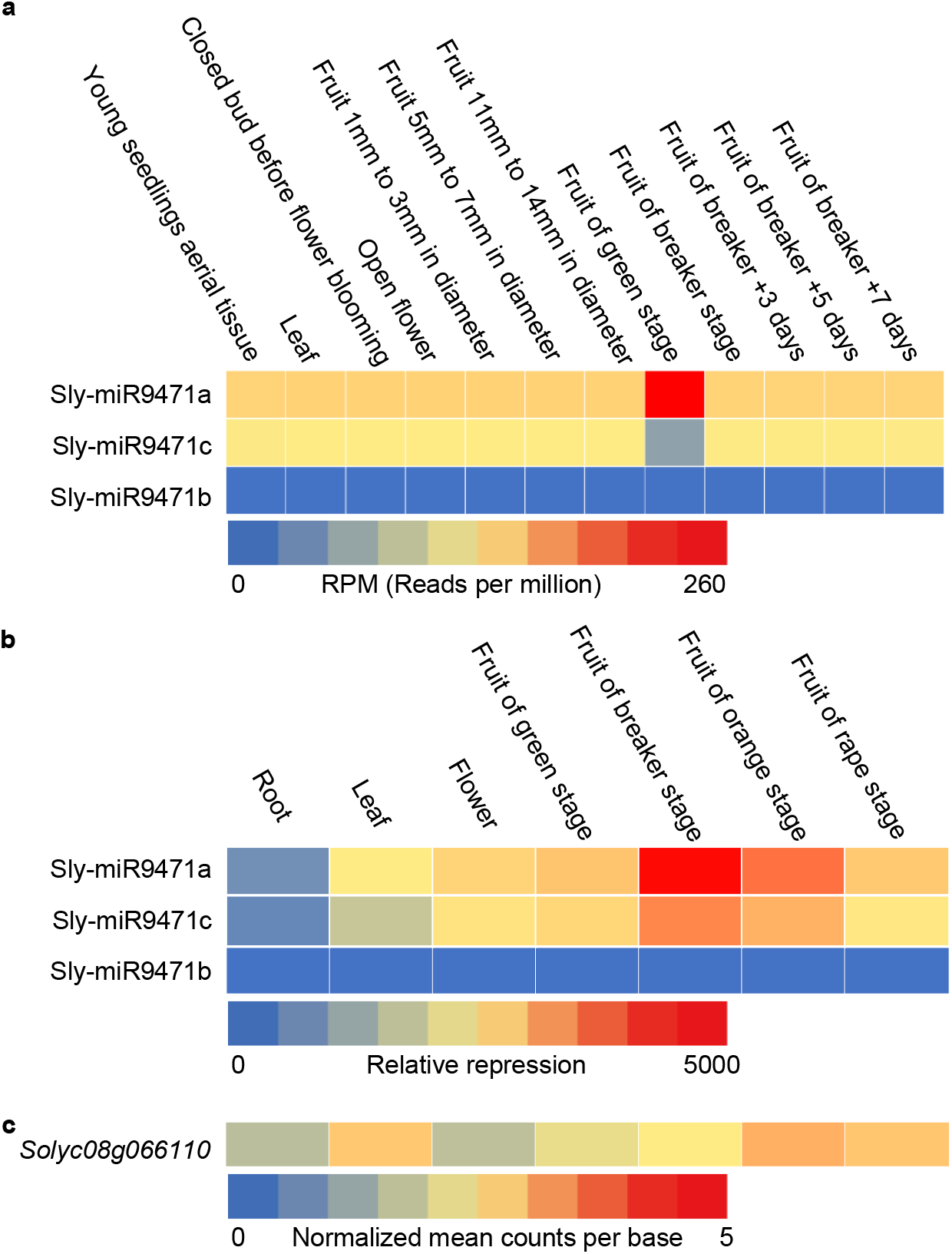
Comparison of the expression pattern of Sly-miR9471 and the host gene *Solyc08g066110* in *S. lycopersicum*. **a**, Expression values of Sly-miR9471a/b/c are determined using sRNA-Seq datasets representing 12 tissues or development stages of the *S. lycopersicum* plants with the accession numbers GSM577998, GSM577999, GSM803579, GSM304986, GSM452712, GSM452713 GSM452714, GSM452715, GSM452716, GSM452717, GSM452718, GSM452719, GSM452720 GSM452721. Colors indicate the expression value in reads per million (RPM). **b**, Relative expression level of Sly-miR9471a/b/c in seven tissues or development stages using RT-qPCR. **c**, Normalized expression of *Solyc08g066110* in the same tissue types as in (**b**). Normalized expression values of *Solyc08g066110* were retrieved from TomExpress (Zouine et al., 2017). Shown are the average expression values in the same tissues types as in (**b**).

**Supplementary Fig. 19.**
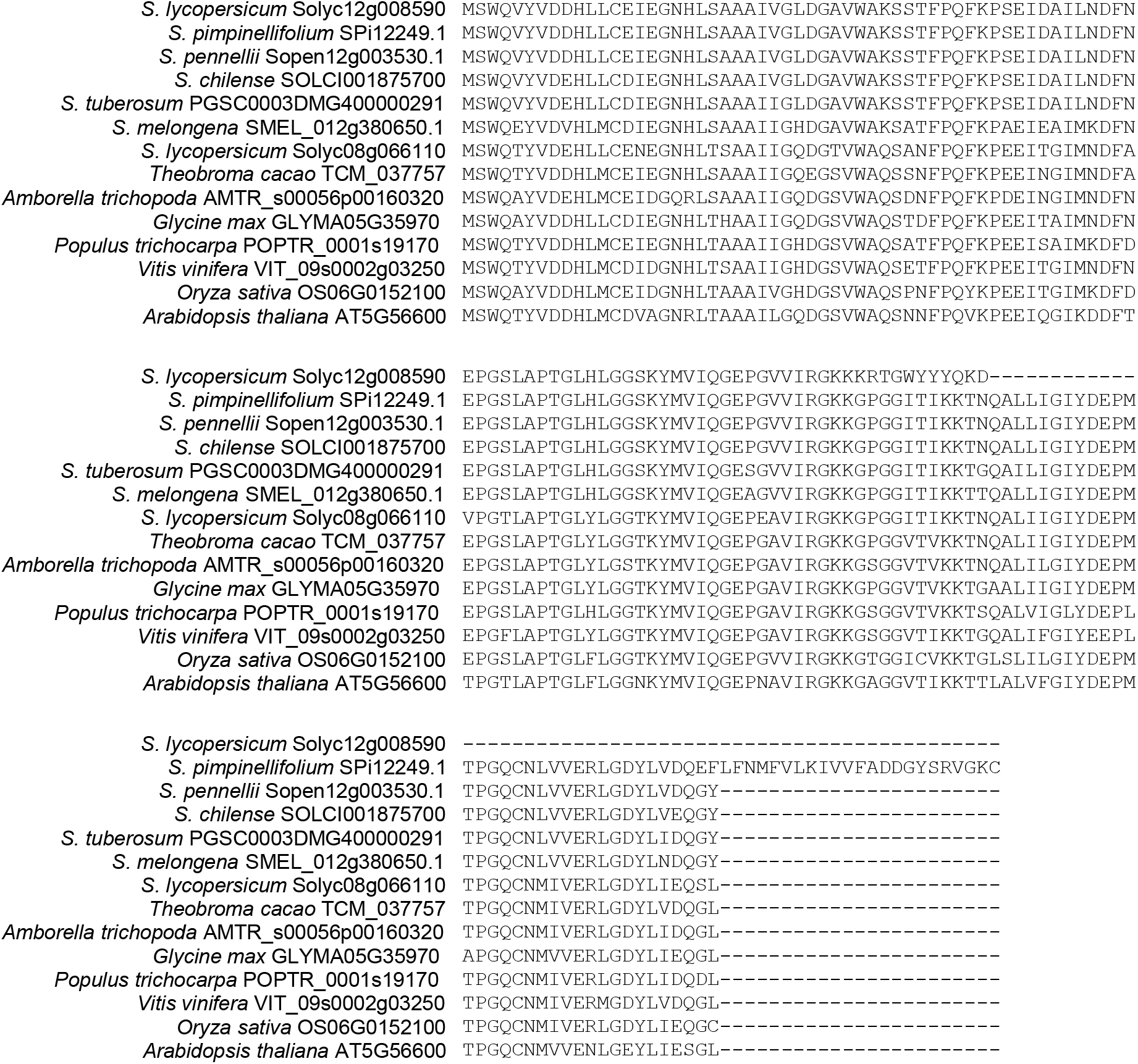
Amino acid alignment of *Solyc12g008590* homologs. Multiple sequence alignment of putative orthologs or paralogs of *Solyc12g008590*. Note that *Solyc12g008590* in *S*. *lycopersicum* and *SPi12249.1* in *S*. *pimpinellifolium* are pseudogenes while other homologs are well conserved.

**Supplementary Fig. 20.**
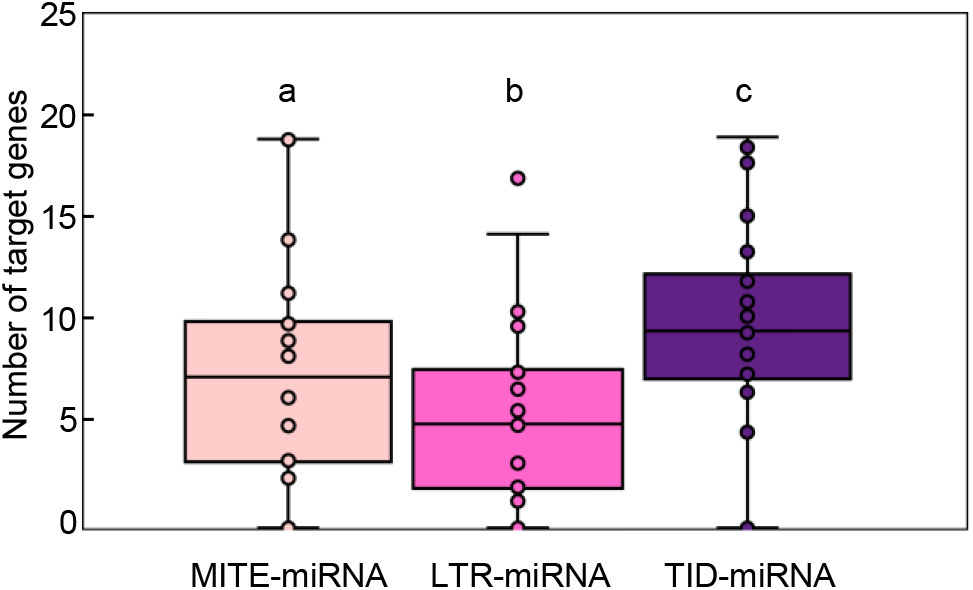
Comparison of the number of predicted target genes of MITE-miRNAs, LTR-miRNAs, and TID-miRNAs. Box plot was used to compare the number of predicted target genes for MITE-miRNAs, LTR-miRNAs, and TID-miRNAs in the examined plant species. The box represents values corresponding to the 25^th^ and 75^th^ percentile. Horizontal line inside the box represents the median. Different letters denote groups with significant differences (*P* < 0.01 by independent samples t-test).

**Supplementary Fig. 21.**
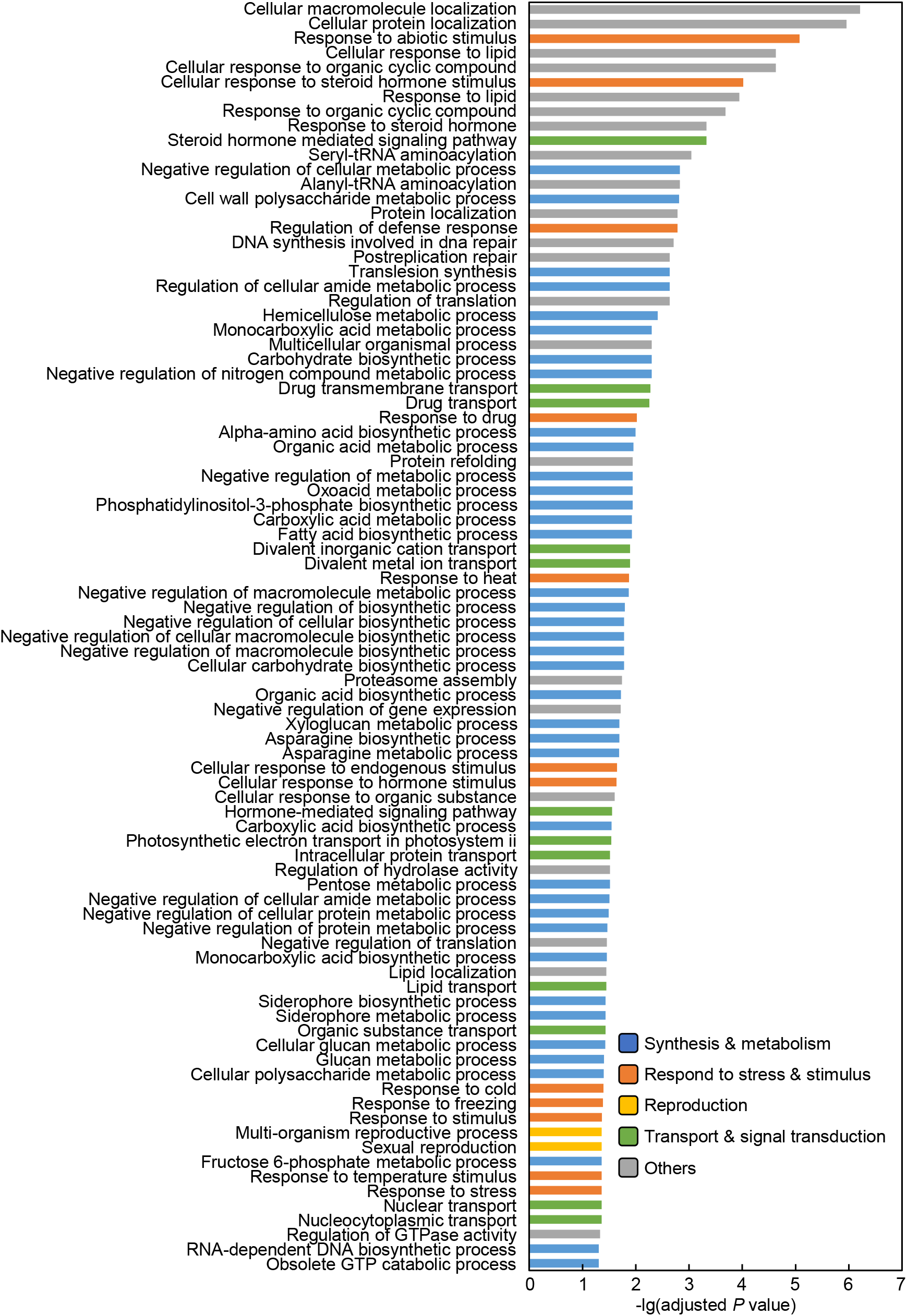
Enriched GO terms related to biological process for the target genes of MITE-miRNAs. Enriched GO terms were identified using Fisher’s exact test with BH adjusted *P* value cutoff at 0.05 was applied.

**Supplementary Fig. 22.**
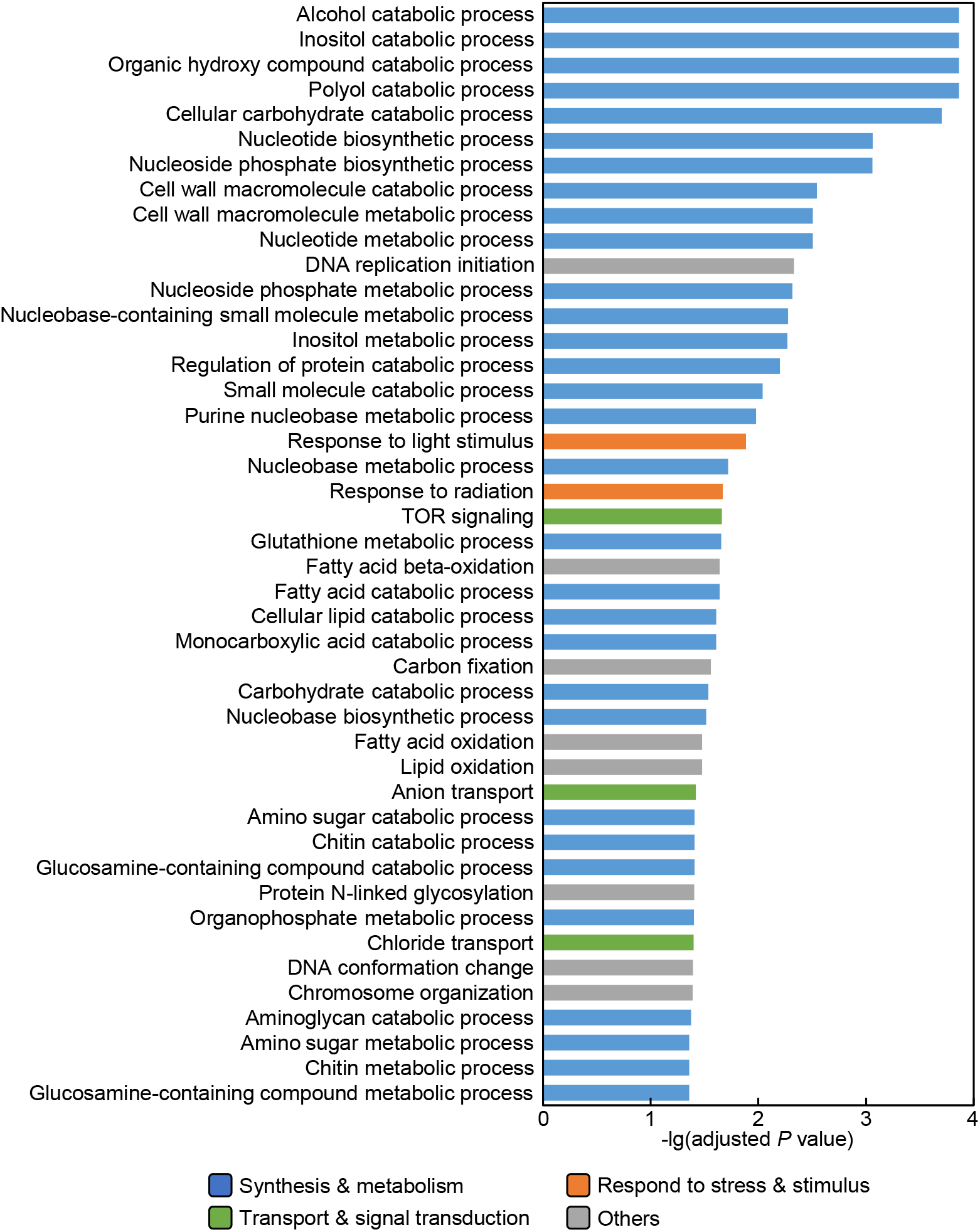
Enriched GO terms related to biological process for the target genes of LTR-miRNAs.

**Supplementary Fig. 23.**
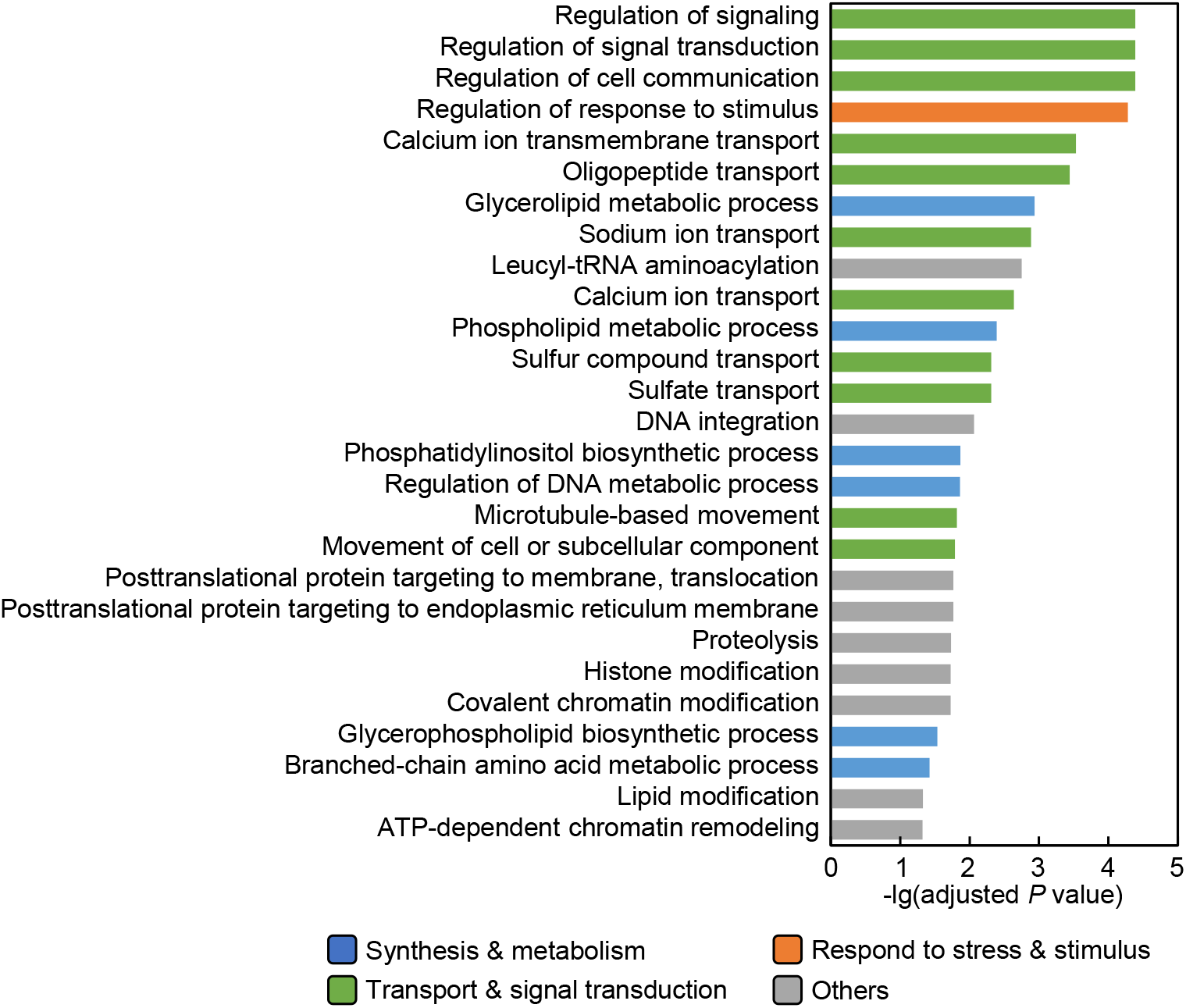
Enriched GO terms related to biological process for the target genes of TID-miRNAs.

**Supplementary Fig. 24.**
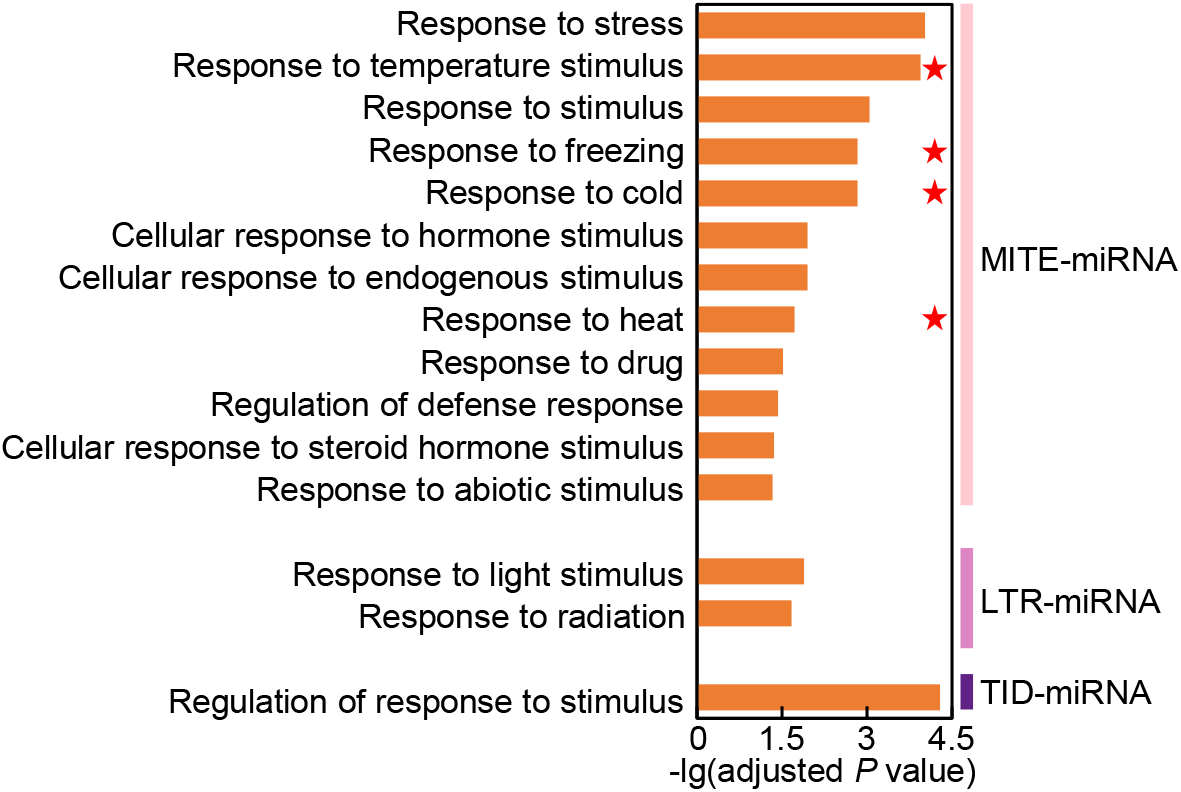
Comparison of enriched GO terms associated with response to stresses and stimuli among the target genes of different types of miRNAs. Significantly enriched GO terms associated with response to stresses and stimuli among the target genes for MITE-miRNAs, LTR-miRNAs, and TID-miRNAs are shown. GO terms related to temperature are marked with red asterisks.

**Supplementary Fig. 25.**
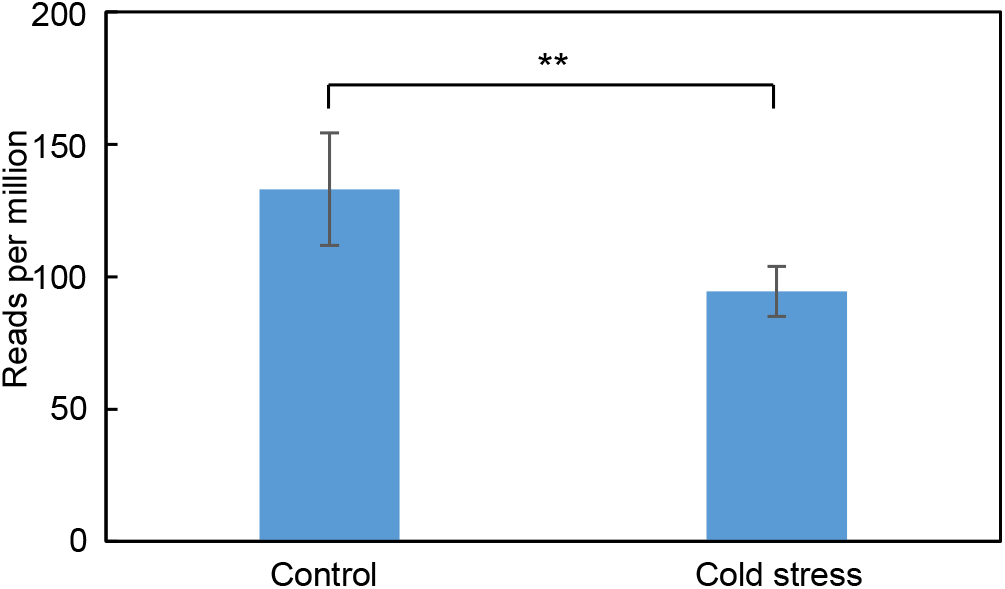
Cold treatment suppresses Osa-miRN2285 expression. Expression of Osa-miRN2285 was compared in cold-stressed and control *O*. *sativa* plants using sRNA-Seq datasets with the accession numbers GSM816687, GSM816688, GSM816689, GSM816694, GSM816695, GSM816730, GSM816731, GSM816732, GSM816739, GSM816740. **, *P* < 0.01 by independent samples t-test.

**Supplementary Fig. 26.**
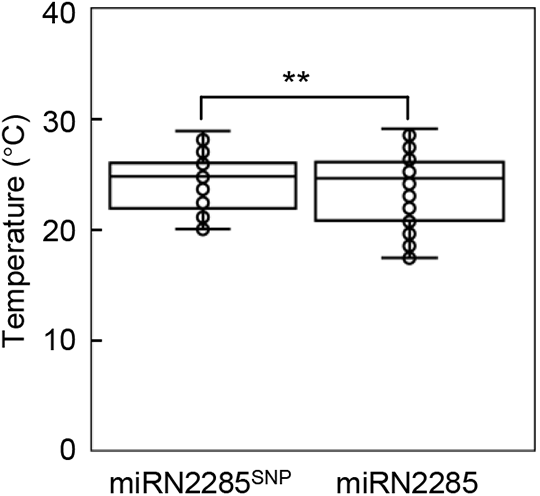
Comparison of summer temperature at the production sites of the miRN2285 and miRN2285^SNP^ type of germplasms. Geographical information for the production sites of the 171 miRN2285^SNP^ type germplasms and the 1,177 miRN2285 type germplasms was retrieved from a public database (Peng et al., 2019). The ten-year (from 2009 to 2018) average summer temperature at the production sites was calculated and compared. **, *P* < 0.01 by independent samples t-test.

